# Scaling, but not instruction tuning, increases large language models’ alignment with language processing in the human brain

**DOI:** 10.1101/2024.08.15.608196

**Authors:** Changjiang Gao, Zhengwu Ma, Jiajun Chen, Ping Li, Shujian Huang, Jixing Li

## Abstract

Transformer-based large language models (LLMs) have significantly advanced our understanding of meaning representation in the human brain. However, increasingly large LLMs have been questioned as valid cognitive models due to their extensive training data and their ability to access context thousands of words long. In this study, we investigated whether instruction tuning, another core technique in recent LLMs beyond mere scaling, can enhance models’ ability to capture linguistic information in the human brain. We evaluated the self-attention of base and fine-tuned LLMs of different sizes against human eye movement and functional magnetic resonance imaging (fMRI) activity patterns during naturalistic reading. We show that scaling has a greater impact than instruction tuning on model-brain alignment, reinforcing the scaling law in brain encoding performance. These finding have significant implications for understanding the cognitive plausibility of LLMs and their role in studying naturalistic language comprehension.

## Introduction

Autoregressive Transformers are increasingly used in cognitive neuroscience for language processing studies, enhancing our understanding of meaning representation and composition in the human language system (Caucheteux & King, 2022; Goldstein et al., 2022; Huth et al., 2016; Schrimpf et al., 2021; Shain et al., 2024; Yu et al., 2024). For instance, Goldstein et al. (2022) found that the probability of words given a context significantly correlates with human brain activity during naturalistic listening, suggesting that language models and the human brain share some computational principles for language processing, such as the “next-word prediction” mechanism (see also Elman, 2004; Hasson et al., 2020). Additionally, pre-trained Transformers are essential for decoding speech or text from neuroimaging data (e.g., Millet et al., 2022; Tang et al., 2023). They provide embeddings for training encoding models that map words to neural data and generate continuations as decoding candidates (Tang et al., 2023). However, those studies mostly adopted smaller pre-trained language models such GPT-2 (Radford et al., 2019) and BERT (Devlin et al., 2019), whereas recent large language models (LLMs) such as GPT-4 (OpenAI et al., 2024) and LLaMA (Touvron et al., 2023) are significantly larger in terms of parameter size and training data. It has been demonstrated that as model size, training dataset and compute budget increase, so does performance on benchmark natural language processing (NLP) tasks, following a power-law scaling law (Henighan et al., 2020; Hestness et al., 2017; Kaplan et al., 2020). These newer LLMs have already been adopted in recent studies to understand language processing in the human brain (e.g., Gao et al., 2024), but whether these LLMs better resemble human language processing remains debated. On the one hand, larger models have been shown to exhibited a stronger correlation with human brain (Antonello et al., 2023; Hong et al., 2024) during language comprehension, mirroring the scaling law in other deep learning contexts. On the other hand, larger models have been questioned as valid cognitive models due to their extensive training data and their ability to access context thousands of words long, which far exceeds human capabilities. Research has shown that surprisal from larger transformer-based language models provide a poorer fit to human reading times (Oh & Schuler, 2023b), and that language model with the lowest perplexity may not result in best model fits to human reading times (Kuribayashi et al., 2021; Oh & Schuler, 2023a). Additionally, limiting the context access of language models can improve their simulation of language processing in humans (Kuribayashi et al., 2022; Yu et al., 2024).

In addition to scaling, fine-tuning LLMs has been shown to improve performance on NLP tasks and enhance generalization to new tasks (Chung et al., 2024; Ouyang et al., 2022; Sanh et al., 2022; Wei et al., 2021). For instance, Ouyang et al. (2022) fine-tuned GPT-3 of varying sizes using reinforcement learning from human feedback (RLHF; Christiano et al., 2017; Stiennon et al., 2020), and showed that the fine-tuned models with only 1.3B parameters were more aligned with human preferences than the 175B base GPT-3. Recent reasoning LLMs such as DeepSeek-R1 (DeepSeek-AI et al., 2024) —which integrates chain-of-thought reasoning with reinforcement learning during fine-tuning—achieve state-of-the-art performance while utilizing similar or fewer activated parameters than existing open-source LLMs. The superior performance of these fine-tuned LLMs over base LLMs on NLP tasks raises the question of whether scaling or fine-tuning have more impact on the models’ brain encoding performance.

In this work, we systematically compared the self-attention of base and fine-tuned LLMs of varying sizes against human eye movement and functional magnetic resonance imaging (fMRI) activity patterns during naturalistic reading (Li et al., 2022). We show that as model size increases from 774M to 65B, the alignment with human eye movement and fMRI activity patterns also significantly improves, adhering to a scaling law (Antonello et al., 2023; Hong et al., 2024). Instruction tuning, on the other hand, does not affect this alignment, consistent with prior findings (Kuribayashi et al., 2024). Model analyses show that base and fine-tuned LLMs diverged the most when instructions were added to the stimuli sentences, suggesting that fine-tuned LLMs are sensitive to instructions in ways that naturalistic human language processing may not be.

## Results

### Model performance on the text stimuli

We used the publicly available Reading Brain dataset from OpenNeuro (Li et al., 2022) to investigate the effects of scaling and instruction tuning on the alignment between LLMs and human eye movement and neural data. The dataset includes concurrent eye-tracking and fMRI data collected from 50 native English speakers (25 females, mean age = 22.5 ± 4.1 years) as they read five English STEM articles inside an fMRI scanner. Each article contains an average of 29.6 ± 0.68 sentences, with each sentence comprising approximately 10.33 ± 0.15 words. Participants read each article sentence-by-sentence in a self-paced manner, pressing a response button to advance to the next sentence. We regressed the self-attention of base and fine-tuned LLMs of varying sizes against the eye movement and functional fMRI activity patterns of each sentence (see Fig. 1 for the experimental procedure and the analyses pipeline). The LLMs employed for our study include all GPT-2 models (base, medium, large, xlarge), 4 different sizes of LLaMA (7B, 13B, 30B, and 65B), two fine-tuned versions of LLaMA (Alpaca and Vicuna) in 7B and 13B configurations and two other fine-tuned models Gemma-Instruct 7B and Mistral-Instruct 7B (see Table 1 for the detailed configurations of the LLMs).

**Fig. 1.**
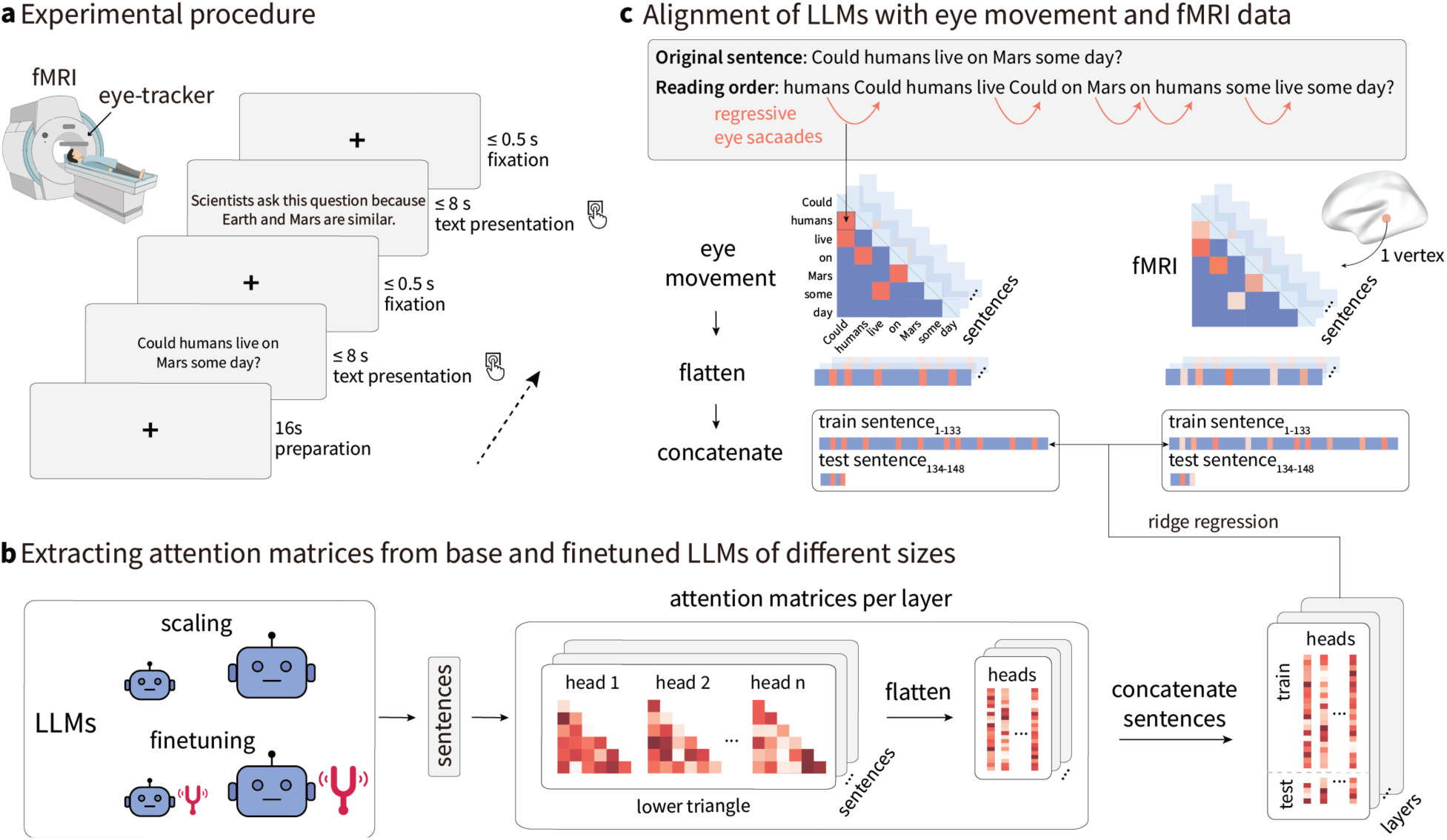
Experimental procedure of the dataset and the analyses pipeline. **a.** Experiment procedure. Participants read five English articles sentence by sentence inside the fMRI scanner with concurrent eye-tracking. **b.** LLMs of different sizes with and without finetuning are employed in the study. **c.** Analysis pipeline. The attention matrices of each layer of the LLMs for each sentence in the experimental stimuli were averaged over attention heads and aligned with eye movement and fMRI activity patterns for each sentence using ridge regression.

**Table 1.**
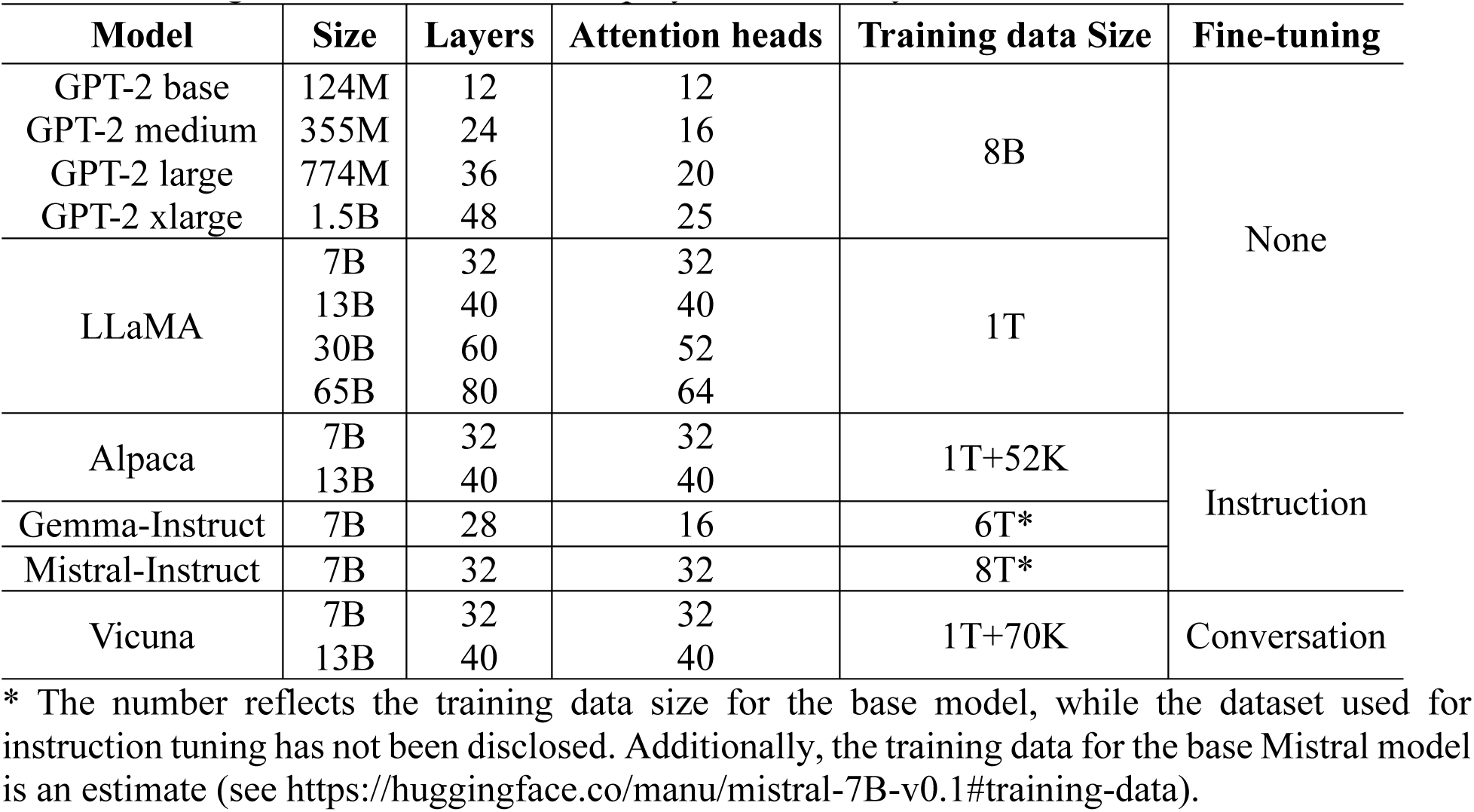
Configurations of the LLMs employed in the study.

Before comparing LLMs with human behavioral and neural patterns, we first evaluated their performance on the experimental stimuli independently. To test how much the LLMs differ in predicting the next word, we calculated the averaged next-word prediction (NWP) loss of all the LLMs on every sentence of our stimuli. The NWP loss exhibited a trend where, for the base models, an increase in model size corresponded to a decrease in mean NWP loss. However, fine-tuned models did not improve performance on NWP for our test stimuli (see Fig. 2a and Supplementary Table 1 for the mean NWP loss for each model. Supplementary Table 2 shows the *t*-test statistics between all model pairs).

**Fig. 2.**
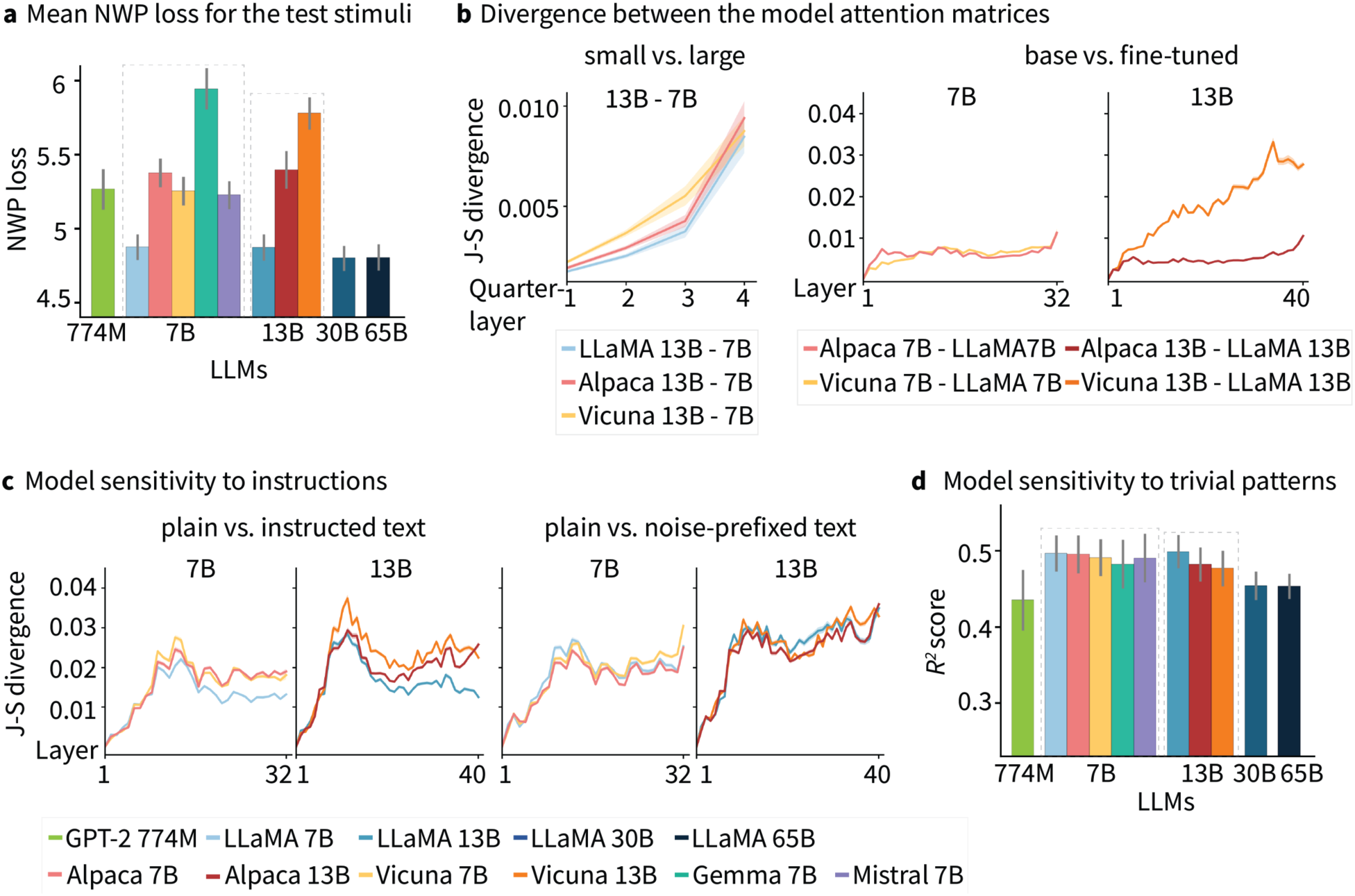
Comparison between the attention matrices of different LLMs. **a,** The mean next-word prediction (NWP) loss of all the LLMs for the test stimuli. **b,** J-S divergence between the attention matrices of different LLMs at each layer or quarter-layer. **c,** The effect of scaling and finetuning on LLMs’ sensitivity to instructions. Shaded regions denote standard deviation. **d,** The effect of scaling and finetuning on LLMs’ sensitivity to trivial patterns between words in a sentence. Error bars denote standard deviation.

### Comparison of model attentions

To examine the effect of scaling and instruction tuning on LLMs’ attention matrices, we calculated the mean Jensen-Shannon (J-S) divergence (*D*_*JS*_) for each pair of LLM’s attention matrices over all attention heads at each model layer. We compared only the LLaMA models and their fine-tuned variants to control for potentially confounding factors such as variations in model architecture and training data. For LLMs with the same number of layers, we computed the *D*_*JS*_layerwise. For LLMs with different numbers of layers, we averaged the attention matrices for every quarter of layers and computed the *D*_*JS*_ for each quarter-layer. Fig. 2b shows the results of the divergence analyses. We observed that for both the base (LLaMA) and fine-tuned models (Alpaca and Vicuna), as the model size increases, the *D*_*JS*_of model attentions linearly increases from the first quarter to the last quarter of the model layers. However, when comparing the base and fine-tuned models of the same sizes, the *D*_*JS*_of model attentions remains small across all layers for most model pairs, except for Vicuna-13B and LLaMA-13B, which exhibit significantly larger divergence, particularly in the higher layers (see Supplementary Table 3 for the detailed *t*-test statistics). Vicuna was fine-tuned using conversational data, incorporating multi-turn dialogues that capture a wide range of conversational contexts (Chiang et al., 2023). As a result, it provides a more natural and context-aware dialogue experience compared to Alpaca, which was fine-tuned on instruction-following examples, leading to strong performance on single-turn tasks (Taori et al., 2023). This distinction may account for the greater divergence observed between Vicuna 13B and LLaMA 13B.

### Sensitivity of model attention to instructions

To confirm that the fine-tuned models exhibit distinct instruction-following behaviors compared to the base models, we analyzed the sensitivity of their attention to instructions. We added two instructions before each sentence in our text stimuli: “Please translate this sentence into German:”, and “Please paraphrase this sentence:”. As a control, we introduced a noise prefix composed of five randomly sampled English words, such as “Cigarette first steel convenience champion.” We then extracted the attention matrices for the original sentence spans and calculate the *D*_*JS*_ of attentions between each model pair layerwise. Our results showed a significantly larger divergence in the attention matrices for the fine-tuned models when processing plain versus instructed texts, for both the 7B and 13B sizes. In contrast, the LLaMA models did not show sensitivity to instructions at either size. No significant difference was found for the *D*_*JS*_of attentions across all layers between the base and fine-tuned models for plain versus noise-prefixed text (see Fig. 2c and Supplementary Table 4 for the detailed *t*-test statistics).

### Sensitivity of model attention to trivial patterns

Prior studies have highlighted certain patterns in LLMs’ attention matrices, such as a tendency to focus on the first word of a sentence, the immediately preceding word (Vig and Belinkov, 2019), or on the word itself (Clark et al., 2019). We consider these tendencies “trivial patterns” because these behaviors are exhibited by all LLMs. As a result, it is not relevant to the effects of scaling or fine-tuning on LLMs’ brain encoding performance, which is the primary focus of this study. To examine how scaling and fine-tuning influence the models’ sensitivity to these trivial patterns, we constructed a binary matrix for each sentence in the test stimuli, marking cells that exhibited these trivial relationships. We then regressed each model’s attention matrix for each sentence at each layer against the corresponding trivial patterns. Our findings showed that for the LLaMA series and their fine-tuned versions, as the model size increases from 7B to 65B, the average regression score for predicting the trivial patterns across layers decreases. No significant differences were observed between the LLaMA models and their fine-tuned versions (see Fig. 2d and Supplementary Table 5-6). Given that similar trivial patterns were not observed in human eye movement data, we believe they do not reflect underlying human cognitive processes. Since the attention weights of larger models display fewer trivial patterns compared to smaller models, this reduced sensitivity may contribute to their greater cognitive plausibility.

### Effects of scaling versus finetuning on model-behavior alignment

Comparisons of LLMs in NWP on our test stimuli indicate an advantage for larger models, suggesting they may achieve better alignment with both behavioral and neural data. To test this hypothesis, we first regressed the attention matrices of the LLMs against the number of regressive eye saccades for all stimuli sentences. We did not include forward saccades, not only due to the unidirectional nature of LLMs but also because regressive saccades may carry more informative value in reading. Regressive saccades occur when readers revisit earlier text, highlighting the importance of previous words in understanding the current word (Liversedge & Findlay, 2000)—similar to how attention weights function in LLMs. We extracted the lower-triangle portions (excluding the diagonal line) of the attention matrix *n*_w*ord*_ × *n*_*word*_ × *n*_*head*_ from all attention heads for every sentence. The attention matrices for all sentences were concatenated to create a regressor with dimensions 7388 × *n*_*head*_ for each layer, where 7388 represents the total number of elements obtained after concatenating the lower triangles of the attention matrices across all sentences in our stimuli. For the human eye saccade data, we constructed matrices for saccade number, 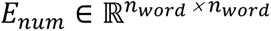, for each sentence. Each cell at row *l* and column *m* in *E*_*num*_ and *E*_*dur*_ represents the number of eye fixation moving from the word in row *l* to the word in column *m*, respectively. We then extracted the lower-triangle parts of the matrices which marks right-to-left eye movement. Similar to the models’ attention matrices, we flattened the regressive eye saccade number matrices for all sentences and concatenated them to create 7388-length vectors for each subject. We then performed ridge regression for each model layer, using the 7388 × *n*_*head*_ regressor to predict each subject’s regressive eye saccade number vectors. The final 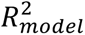 was normalized by the 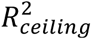, where the ceiling model represents the mean of all subjects’ regressive eye saccade number vectors.

Our findings show that for the LLaMA series, as model size increases from 7B to 65B, the regression scores also increase across layers. The GPT-2 models, which has the smallest parameter size, exhibits the lowest regression scores. In contrast, base and fine-tuned models of the same sizes exhibit no difference in their regression scores when aligned with human eye movement patterns, suggesting that scaling, rather than finetuning, enhances the alignment between LLMs and human reading behaviors. No significant difference was found for the regression scores of controlled models of matching sizes (see Fig. 3a and Supplementary Table 7-8). Notably, the GPT-2 models of varying sizes did not exhibit any significant differences in the fit between these models and the eye-regression patterns. This may be because the size differences among these models are not as substantial as, for example, between 7B and 65B. We further plotted the maximum regression scores from all model layers against different LLMs and the logarithmic scale of parameter size, illustrating a clear scaling law of model-behavior alignment (see Fig. 3a, right panel).

**Fig. 3.**
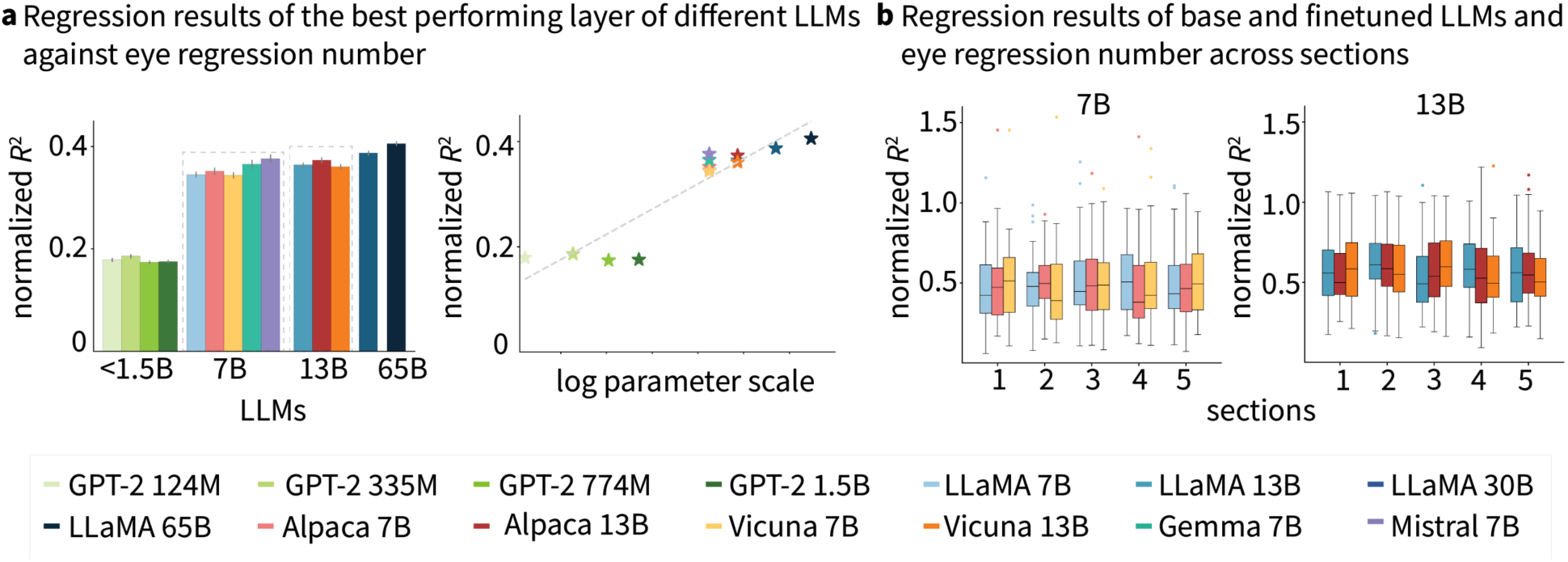
Effects of scaling and finetuning on the alignment between LLMs and human regressive eye saccade patterns during naturalistic reading. **a,** Regression results of the LLMs’ best performing layer on the regressive eye saccade patterns and their results on a logarithmic size scale. Error bars denote standard deviation. **b,** Regression results of different LLMs and regressive eye saccade number patterns across experimental sections.

Given that participants answered 10 comprehension questions after reading each article, there is a possibility that their reading behavior shifted from naturalistic reading to a more focused approach aimed at solving questions as the experiment progressed. This could mean that LLMs with instruction tuning might increasingly align with human behavior later in the experiment. To test this hypothesis, we performed the same regression analyses separately for each section of the experiment. Our results revealed no significant difference in the regression scores for base and fine-tuned LLMs over time, suggesting that human reading behaviors during naturalistic reading are not influenced by the subsequent comprehension questions (see Fig. 3b). Supplementary Table 9 lists the *F* statistics from one-way analysis of variance (ANOVA) for each base and fine-tuned LLM across the 5 experimental sections.

### Effects of scaling versus finetuning on model-brain alignment

We next conducted a ridge regression using the attention matrix 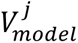 at each layer *j* from each LLM to predict each voxel’s BOLD vector 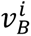 in the whole brain for each subject (see “Alignment between LLMs and fMRI data” in Methods for details). As shown in Fig. 4a, the average prediction performance across subjects from the best performing layer of each LLM increases with model size. In contrast, base and fine-tuned LLMs of the same sizes did not show differences in their average prediction performance (see Supplementary Table 10-11). We also plotted the normalized correlation coefficients from the best-performing layer of each model on a logarithmic scale, demonstrating a clear scaling effect where larger models better explained the fMRI activity patterns during naturalistic reading. Fig. 4b presents significant brain clusters identified when contrasting the prediction performance (Pearson’s r maps) of larger and smaller LLMs. The results show that larger LLMs consistently exhibited significantly more activation in a bilateral temporal-parietal network compared to their smaller counterparts. Although the effect size as measured by the Cohen’s d is larger in the left hemisphere than the right hemisphere (see Table 2). Additionally, we compared the regression scores of base and fine-tuned models of same sizes, yet no significant brain cluster has been observed.

**Fig. 4.**
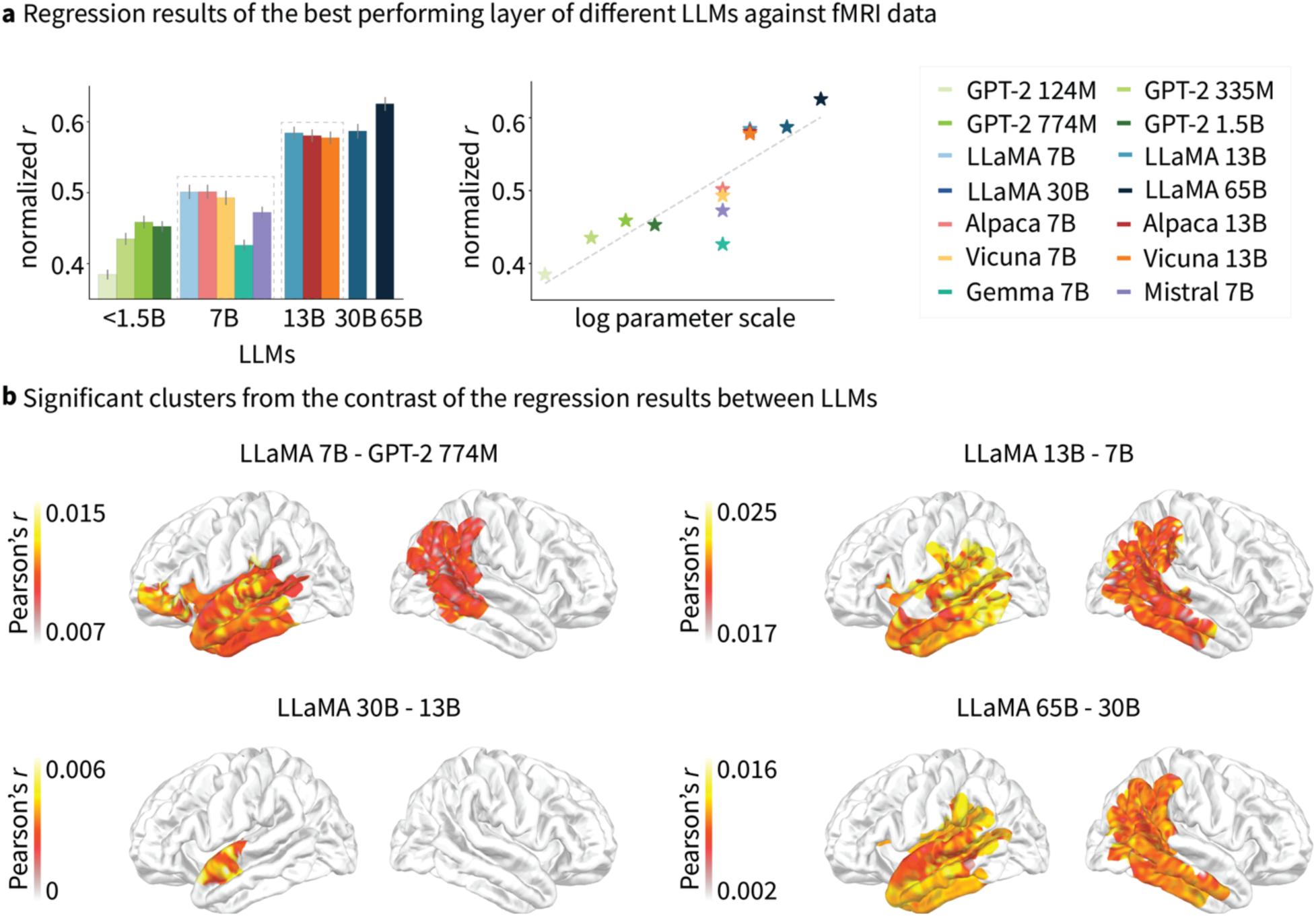
Effects of scaling and finetuning on the alignment between LLMs and human fMRI activity patterns during naturalistic reading. **a,** Regression results of the LLMs’ best-performing layer on the fMRI activity patterns and their results on a logarithmic size scale. Error bars denote standard deviation. **b,** Significant brain clusters from the contrast of the correlation coefficients of different LLMs with smaller and larger sizes.

**Table 2.** Summary statistics for the significant brain clusters in the left and right hemisphere from the contrast of the correlation coefficients of model pairs from the ridge regression analyses.

| Model 1 | Model 2 | Left hemisphere |  |  | Right hemisphere |  |  |
| --- | --- | --- | --- | --- | --- | --- | --- |
| | | N vertices | $p$ | Cohen's d | N vertices | $p$ | Cohen's d |
| LLaMA 7B | GPT-2 large | 1133 | 0.006 | 7.303 | 517 | 0.007 | 3.691 |
| LLaMA 13B | LLaMA 7B | 1096 | <0.001 | 18.005 | 622 | <0.001 | 7.670 |
| LLaMA 30B | LLaMA 13B | 69 | <0.1 | 2.159 | \ | \ | \ |
| LLaMA 65B | LLaMA 30B | 1734 | 0.007 | 7.149 | 650 | 0.008 | 3.709 |

### Expanding the analysis to different datasets

To verify whether our findings can generalize to a broader spectrum of human language processing, we performed the same analysis on a fMRI dataset collected while participants listened to a 20-minute Chinese audiobook in the scanner. We regressed the attention weights of the base and fine-tuned LLaMA3 8B, LLaMA3 70B, LLaMA3-Instruct 8B and LLaMA3-Instruct 70B models against the fMRI data matrices at the paragraph level (See “Additional results from fMRI data of naturalistic listening” in the Supplementary information). We used the LLaMA3 models for their better performance in Chinese. This analysis extends beyond sentence-level comprehension to discourse-level processing and introduces both a different modality (listening vs. reading) and a different language (Chinese vs. English). Our findings remained consistent: Model scaling had a significant effect on model-brain alignment, while fine-tuned and base models of the same size showed no difference in brain encoding performance (see Supplementary Fig. 1a and Supplementary Table 12-14).

Additionally, we regressed the predictions from the base and fine-tuned LLaMA3 8B and LLaMA3 70B models against the fMRI data collected while participants answered multiple-choice comprehension questions about the preceding listening session (See “Additional results from fMRI data of naturalistic listening” in the Supplementary information). Our results showed LLaMA3 8B exhibited a significantly higher regression score (mean=0.219±0.004) compared to LLaMA3-Instruct 8B (mean=0.211±0.004, t=58.073, p<0.001); LLaMA3-Instruct 13B showed a higher mean regression score (mean=0.259±0.005) across participants compared to the LLaMA3 13B (mean=0.243±0.005, t=80.528, p<0.001), but no significant brain cluster has been found between the contrast of the two 13B models’ R^2^ maps (see Supplementary Fig. 1b and Supplementary Table 15-17).

## Discussion

Scaling and finetuning are two key factors behind the enhancements of recent LLMs compared to their predecessors, such as BERT (Devlin et al., 2019) and GPT-2 (Radford et al., 2019). While prior work suggests that the scaling law also applies to LLMs’ brain encoding performance with extensive fMRI data during naturalistic language comprehension (Antonello et al., 2023), the efficacy of scaling in model-brain alignment with shorter texts remains uncertain. As it has been shown that GPT-2 predicts more variance relative to the ceiling in some neuroimaging datasets (Fedorenko et al., 2016; Pereira et al., 2018)—defined as the mean of the neural responses from all participants in these datasets, smaller models may have reached their performance limits on next-word prediction for simpler texts. Moreover, current LLMs far exceed human capabilities in terms of data input during training and memory resources for accessing contextual information during comprehension. It has been argued that larger models increasingly diverge from human language processing patterns (Kuribayashi et al., 2022). In this study, we evaluated the alignment between LLMs of varying sizes and human eye movement and fMRI activity patterns during naturalistic reading. Despite using experimental stimuli and fMRI data much smaller in size compared to previous studies (Antonello et al., 2023), we observed consistent improvements in alignment as model size increased from 774M to 65B, without any apparent diminishing returns. Similar results have also been reported for ECoG data during naturalistic listening (Hong et al., 2024). This suggests that the scaling law of model-brain alignment holds even with shorter text stimuli and smaller fMRI data.

While the largest LLMs today still do not match the human brain in terms of synapse count, training and operating such large LLMs pose significant computational challenges, especially in academic settings with limited computing budgets. Finetuning LLMs with instructions offers a viable approach to enhance the performance and usability of pre-trained language models without expanding their size (Chung et al., 2024; Ouyang et al., 2022; Sanh et al., 2022; Wei et al., 2021). Ouyang et al. (2022) noted that the typical “next-token prediction” training objective of language models often diverges from user intentions, leading to outputs that are less aligned with user preferences. Although there is ample evidence that human language processing involves “next-word prediction” (Goldstein et al., 2022; Hasson et al., 2020; Ryskin & Nieuwland, 2023), research also showed that finetuning language models for tasks like narrative summarization can enhance model-brain alignment, especially in understanding characters, emotions, and motions (Aw & Toneva, 2022). It is possible that “instruction-following” plays a role in human language learning and that fine-tuned models might contain richer discourse and pragmatic information beyond basic meaning representation.

However, our regression results with human behavioral and neural patterns did not reveal any significant improvement in alignment for fine-tuned LLMs compared to base models of the same sizes. We examined whether fine-tuned models exhibited better alignment to eye movement patterns as participants completed more comprehension questions over time, but no significant differences were found in the regression scores. We also examined predictions from the fine-tuned LLaMA3 7B and LLaMA3 70B models against the fMRI data collected while participants answered multiple-choice comprehension questions about the preceding listening session, yet we still did not find a consistent advantage of the fine-tuned model on model-brain alignment. Our results therefore highlight the greater impact of scaling over fine-tuning in model-brain alignment, contributing to the existing literature on the scaling law in brain encoding performance (Antonello et al., 2023; Hong et al., 2024; Schrimpf et al., 2021). Similar findings have been reported by Kuribayashi et al. (2024), who demonstrated that instruction-tuned and prompted LLMs do not provide better estimates than base LLMs when simulating human reading behavior. However, it is possible that LLMs using different fine-tuning techniques may exhibit a positive effect. Here we examined two additional fine-tuned models (Gemma-Instruct and Mistral-Instruct) and did not find any improvement over the base LLMs, but Kuribayashi et al. (2024) reported that Falcon instruction-tuned LLMs, which utilize a supervised tuning approach different from RLHF, showed a moderate positive effect in simulating human reading data. Future research should further explore the impact of fine-tuning techniques on the cognitive plausibility of instruction tuning.

Our findings that scaling has a larger impact than fine-tuning on model-behavior and model-brain alignments are particularly relevant in the current landscape, where reasoning LLMs such as DeepSeek-R1 (DeepSeek-AI et al., 2024) showed superior performance with similar or fewer activated parameters than existing open-source LLMs. We acknowledge that caution is needed when interpreting these results. Since instruction tuning effectively realigned the model weights in response to instructions, these realigned model weights may better fit brain activity patterns where participants performed tasks aligned with the instruction-following nature of the fine-tuning process. However, due to the lack of such openly available neuroimaging datasets, we cannot evaluated the fine-tuned LLMs on these task-specific brain data.

In summary, we compared base and fine-tuned LLMs of varying sizes against human eye movement and fMRI activity patterns during naturalistic reading. Our results highlighted a significant impact of scaling on model-brain alignment, whereas instruction tuning showed no such effect. These results serve as a reference for researchers selecting LLMs for brain encoding and decoding studies. One limitation of the study is the lack of comparisons between base and fine-tuned LLMs against human behavioral and neural data during experimental tasks with instructions. This gap leaves the potential for future research to explore the impact of instruction tuning on model-brain alignment in controlled experimental settings.

## Methods

### Eye-tracking and fMRI data

We used the openly available Reading Brain dataset (Li et al., 2022) on OpenNeuro. This dataset includes concurrent eye-tracking and fMRI data collected from 52 native English speakers (27 females, mean age = 22.8 ± 4.7 years) as they read five English STEM articles inside an fMRI scanner. 2 subjects’ (sub-21 and sub-52) data were subsequently removed due to preprocessing errors, resulting in 50 subjects’ (25 females, mean age = 22.5 ± 4.1 years) data in total. The articles were constructed using materials from established sources, including the NASA science website, the GPS.gov website (http://www.gps.gov), and Wikipedia. These texts underwent an extensive revision process to ensure content accuracy and stylistic consistency (see Follmer et al., 2018). Each article contains an average of 29.6 ± 0.68 sentences, with each sentence comprising approximately 10.33 ± 0.15 words. Participants read each article sentence-by-sentence in a self-paced manner, pressing a response button to advance to the next sentence. If there was no response within 8000 ms, the screen would automatically progress to the next sentence. The sequence in which the five texts were presented was randomized across participants to control for potential order effects. At the end of each article, participants answered 10 multiple-choice questions to ensure their comprehension. The whole experiment, including preparation time, lasted for about one hour (see Fig. 1a for the experimental procedure).

All imaging and eye-tracking data were acquired at the Center for NMR Research at the Pennsylvania State University Hershey Medical Center in Hershey, Pennsylvania. The anatomical scans were acquired using a Magnetization Prepared RApid Gradient-Echo (MP-RAGE) pulse sequence with T1 weighted contrast (176 ascending sagittal slices with A/P phase encoding direction; voxel size = 1mm isotropic; FOV = 256 mm; TR = 1540 ms; TE = 2.34 ms; acquisition time = 216 s; flip angle = 9°; GRAPPA in-plane acceleration factor = 2; brain coverage is complete for cerebrum, cerebellum and brain stem). The functional scans were acquired using T2* weighted echo planar sequence images (30 interleaved axial slices with A/P phase encoding direction; voxel size = 3 mm × 3mm × 4 mm; FOV = 240 mm; TR = 400 ms; TE = 30 ms; acquisition time varied on the speed of self-paced reading, maximal 5.1 minutes per run; multiband acceleration factor for parallel slice acquisition = 6; flip angle = 35°; where the brain coverage missed the top of the parietal lobe and the lower end of the cerebellum). A pair of spin echo sequence images with A/P and P/A phase encoding direction (30 axial interleaved slices; voxel size=3mm× 3mm× 4mm; FOV=240mm; TR=3000ms; TE=51.2 ms; flip angle = 90°) were collected to calculate distortion correction for the multiband sequences (Glasser et al., 2013). fMRI preprocessing of the was conducted using fMRIPrep (v25.0.0) with all default parameters. Anatomical images were corrected for intensity non-uniformity (N4BiasFieldCorrection), skull-stripped using ANTs-based extraction (OASIS30ANTs template), and segmented into tissue classes using FSL’s ‘fast’. Brain surfaces were reconstructed using ‘recon-all’ (FreeSurfer 7.3.2). The T1-weighted images were normalized to MNI152NLin2009cAsym:res-2 space via nonlinear registration (ANTs). For each BOLD run, head-motion parameters with respect to the BOLD reference (transformation matrices, and six corresponding rotation and translation parameters) are estimated before any spatiotemporal filtering using ‘mcflirt’ (FSL 5.0.9) and slice timing correction was applied using 3dTshift (AFNI 20160207). The BOLD reference was then co-registered to the T1w reference using bbregister which implements boundary-based registration. Co-registration was configured with six degrees of freedom. No susceptibility distortion correction was applied. Confound regressors included motion parameters (and their derivatives/quadratics), framewise displacement (FD), DVARS, global signals, and t/aCompCor components computed from white matter and CSF after high-pass filtering (128s cutoff). Volumes exceeding FD>0.5 mm or standardized DVARS>1.5 were flagged as motion outliers. Final resampling to MNI space and fsaverage5 surface was performed in a single interpolation step using antsApplyTransforms and mri_vol2surf.

Participants’ eye movements were simultaneously recorded using an MRI-compatible EyeLink 1000 Plus eye tracker (SR Research, 2016) with a sampling rate of 1000 Hz. The eye tracker was mounted at the rear end of the scanner bore and captured eye movements via a reflective mirror positioned above the MRI’s head coil. This setup included the following parameters: distance between the reader’s eyes and the presenting screen = 143 cm; distance between the camera and the participant’s eyes via the reflective mirror was 120 cm; presenting screen size = 35.7 cm × 57.2 cm; average word length on the screen = 3.08 cm; average distance between words = 0.95 cm. On average, a reader’s visual angle when fixating on a word is 1°14¢. Recording was performed monocularly from the right eye, and the participant’s head was stabilized using the head coil. A 13-point calibration routine was conducted at the beginning of the experiment, followed by a validation process during the first run. For subsequent runs, validation checks were performed regularly, and if the error exceeded 1°, recalibration was conducted. Participants initially viewed a fixation cross for 500 ms on the left side of the screen before each sentence was displayed, helping them to anticipate the text presentation. The initial fixation cross lasted 6000 ms to allow participants ample time to prepare and for the blood-oxygen-level-dependent (BOLD) signal to stabilize. Subsequent fixation crosses between sentences were displayed for 500 ms each. Due to fixation drifting caused by the declining accuracy of calibration over time, we manually adjusted fixations falling outside the range of predefined target regions where sentences are presented using the Data Viewer^TM^ software from SR Research (SR Research, 2016). Rather than using an auto-adjustment that aligns all fixations to a single horizontal line, we applied trial-by-trial corrections only along the y-axis. This approach preserved the original fixation patterns of the readers. Approximately 15% of the data required this type of manual correction.

### LLMs

To investigate the effects of scaling and instruction tuning on the alignment of LLMs with human behavior and neural data, we utilized the open-source LLaMA model (Touvron et al., 2023) and its instruction-tuned variants, Alpaca (Taori et al., 2023) and Vicuna (Chiang et al., 2023), available in various sizes. LLaMA is a series of pre-trained causal language models trained on over 1 trillion publicly accessible text tokens, primarily in English. It achieved state-of-the-art performance on most LLM benchmarks (Touvron et al., 2023). We employed all four sizes of LLaMA: 7B, 13B, 30B, and 65B. Additionally, we included all the GPT-2 models (Radford et al., 2019) to represent smaller pre-trained language models (base:124M, medium:355M, large:774M, xlarge:1.5B) as well as two other fine-tuned models, Gemma-Instruct 7B (Gemma Team, 2024) and Mistral-Instruct 7B (v.03; Jiang et al., 2023), for comparison with LLaMA 7B.

Alpaca (Taori et al., 2023) was fine-tuned from the 7B LLaMA model and was trained on 52K English instruction-following demonstrations generated by GPT-3 (Brown et al., 2020) using the self-instruct method (Wang et al., 2023). We also developed a 13B version of Alpaca using the same training data and strategy. Our 13B Alpaca model achieved accuracy scores of 43.9 and 46.0 on the MMLU dataset (Hendrycks et al., 2020) in zero-shot and one-shot settings, respectively, outperforming the original 7B model’s scores of 40.9 and 39.2. Vicuna versions 7B and 13B (Chiang et al., 2023) were fine-tuned from the respective 7B and 13B LLaMA models, using 70K user-shared conversations with ChatGPT (OpenAI et al., 2024). This dataset includes instruction and in-context learning samples across multiple languages. Gemma-Instruct 7B was fine-tuned on a mix of synthetic and human-generated prompt-response pairs (Gemma Team, 2024), and Mistral-Instruct 7B was fine-tuned on publicly available instruction datasets from the Hugging Face repository (Jiang et al., 2023).

### Comparison of next-word prediction loss

To examine the effects of scaling and finetuning model’s performance in next-word prediction, we calculated the mean NWP loss (the negative log-likelihood loss normalized by sequence lengths) of the models employed in this study on every sentence of the articles in the Reading Brain dataset. Since LLMs use subword tokenization (Kudo and Richardson, 2018), we aligned subwords to words by summing over the “to” tokens and averaging over the “from” tokens in a split word, as suggested by Clark et al. (2019) and Manning et al. (2020). For example, suppose the phrase “delicious cupcake” is tokenized as “del icious cup cakes”, the attention score from “cupcake” to “delicious” is the sum of the attention scores from “del” to “cup” and “cake” and “icious” to “cup” and “cake”, divided by 2 as there are to “to” tokens (“cup” and “cake”). We also removed the special tokens “<s>” at sentence beginning positions. The losses for all sentences were z-scored model-wise and the contrasts of the z-scored losses for two models (e.g., LLaMA 7B vs. Alpaca 7B) were tested using a two-sample two-tailed related *t*-test. The false discovery rate (FDR) was applied to correct for multiple comparisons across layers.

### Comparison of model attentions

The self-attention matrices of different LLMs given the same input were compared using their mean Jensen-Shannon (J-S) divergence across all layers. For every sentence in our stimuli, we extracted the attention matrices *A* and *B* from one attention head and one layer of two target LLMs 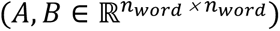, and their J-S divergence *D*_*JS*_(*A*, *B*) is computed as 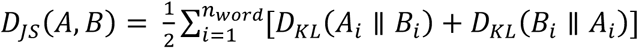,where *A*_*i*_ and *B*_*i*_ are the *i* -th rows in the two matrices and *D*_*KL*_ is the Kullback–Leibler (K-L) divergence (Kullback & Leibler, 1951). The attention matrices were normalized such that each row sums to 1, and the final *D*_*JS*_ for each layer was averaged across attention heads. We aligned subword tokenization to words using the previously described methods for calculating NWP loss. For models with different numbers of layers, we divided their layers into four quarters and averaged the *D*_*JS*_ quarter-wise. We compared each model-pair’s *D*_*JS*_ for each layer or each quarter-layer using a two-sided two-sample related *t*-test with FDR correction.

### Model sensitivity to instructions

We compared the models’ attention matrices for each stimuli sentence when prefixed with two instructions: “Please translate this sentence into German:”, and “Please paraphrase this sentence:”. As a control, we introduced a noise prefix composed of five randomly sampled English words, such as “Cigarette first steel convenience champion.” We then extracted the attention matrices for the original sentence spans. We calculated the *D*_*JS*_ between the prefixed and original sentences across different models to assess each model’s sensitivity to instructions.

### Model sensitivity to trivial patterns

Prior studies have highlighted certain trivial patterns in the attention matrices within a given context, such as a tendency to focus on the first word of a sentence, the immediately preceding word (Vig & Belinkov, 2019), or the word itself (Clark et al., 2019). We consider the model’s tendencies to attend to the immediately preceding or current word “trivial patterns” because these behaviours are exhibited by all LLMs. As a result, it is not relevant to the effects of scaling or fine-tuning on LLMs’ brain encoding performance, which is the primary focus of this study. To examine whether scaling and finetuning will change the models’ reliance on these trivial patterns, we constructed a binary matrix for each sentence in the test stimuli, marking cells that exhibit these trivial attention relationships with a 1. We then flattened the lower-triangle parts of these matrices to create trivial pattern vectors. Subsequently, we performed ridge regressions using each model’s attention vectors for each sentence at each quarter-layer to predict the corresponding trivial pattern vectors. The resulting regression scores were averaged across model layers and were z-scored and assessed for statistical significance using two-tailed one-sample *t*-tests with FDR corrections. We subtracted these patterns from all the attention matrices of the LLMs for the following ridge regression analyses. Given that similar patterns were not observed in human eye movement data, we believe they do not reflect underlying human cognitive processes.

We also examined results based on the original attention matrices (without subtracting the trivial patterns) from each LLM and observed a similar scaling effect, with larger models exhibiting higher model-brain alignment within a bilateral temporal-parietal network. However, no significant brain clusters were observed for the contrasts between the mid-sized LLaMA pairs 13B vs. 7B and 30B vs. 13B (see Supplementary Fig. 2 and Supplementary Table 18–19). This may be due to smaller and mid-sized models being more susceptible to capturing trivial patterns, leading to similar brain responses.

### Alignment between LLMs and eye movement

We input each sentence into the LLMs individually, consistent with how sentences were presented separately to participants on the screen during fMRI scanning. Furthermore, regressive eye saccade information was available only at the sentence level. Since our auto-regressive LLMs utilize right-to-left self-attention, we extracted the lower-triangle portions of the attention matrix from each layer and each attention head for every sentence. These matrices were flattened and concatenated to form the attention vector 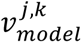 for all sentences at head *k* in layer *j*. We stacked these vectors along the attention heads to create a matrix 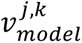 for the *j*-th layer. For the human eye saccade data, we constructed matrices for saccade number 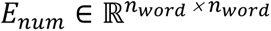, for each sentence. Each cell at row *l* and column *m* in *E*_*num*_ represents the number of times of eye fixation moving from the word in row *l* to the word in column *m*, respectively. We then extracted the lower-triangle parts of the matrices which marks right-to-left regression. Like the models’ attention matrices, we flattened the regressive eye saccade number matrices for all sentences and concatenated them to get the regressive eye saccade number vector 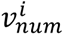 for each subject *i*. We then conducted a ridge regression using the model attention matrix 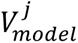 at each layer *j* to predict each subject’s regressive eye saccade number vector. The final dimensionality of the dependent variable X is 7388 × N_att_head_, where 7388 represents the length of the concatenated vector formed by flattening the lower-triangular part of the attention matrix for each sentence at each layer, and N is the number of attention heads at that layer. We did not average across attention heads to obtain a single attention matrix per sentence, as each head is known to capture distinct relationships among words in a sentence (Manning et al., 2020). The dependent variable (y) is a 7388-dimensional vector, where 7388 again reflects the length of the concatenated and flattened lower-triangular part of the regressive eye saccade matrix for each sentence.

We used ridge regression instead of ordinary least squares (OLS) regression as most models have 32 attention heads, with the maximum being 64, we believe that applying ridge regression is preferable to mitigate collinearity among the regressors and enhance prediction accuracy. The penalty regularization parameter was kept as the default value of 1. The final 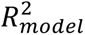 was normalized by the 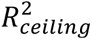, where the ceiling model represents the mean of all subjects’ regressive eye saccade number vectors. At the group level, the significance of the contrast of the regression performance 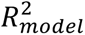 for every model pair at every layer was examined using a two-tailed one-sample related *t*-test, with FDR corrections for multiple comparisons across layers.

To test this hypothesis that participants’ reading behavior shifted from naturalistic reading to a more focused approach aimed at solving questions as the experiment progressed, we performed the same regression analysis separately for each section of the experiment. We then compared the regression scores of each LLM across different times using analysis of variance (ANOVA) to assess changes in model fit over the course of the experiment.

### Alignment between LLMs and fMRI data

For each voxel of the fMRI data for each subject, we constructed a BOLD matrix 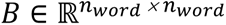 for each sentence. The value at row *l* and column *m* in *B* represents the sum of the BOLD signals at the timepoints where the eye fixation moves from the word in row *l* to the word in column *m*. We extracted the lower-triangle parts of the *B* matrices (excluding the diagonal line) for all sentences and concatenated them to form the BOLD vector 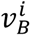 for each voxel of each subject *i*. Next, we conducted a ridge regression using each subject’s regressive eye saccade vectors 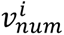 to predict each subject’s BOLD vector 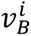 at each voxel. We then performed ridge regressions using the attention matrix 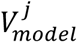 at each layer *j* from each LLM to predict each voxel’s BOLD vector 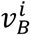 in the whole brain for each subject (see Fig. 1). The dimensionality of the independent variable is 7388 × N_att_head_, where 7388 represents the length of the concatenated vector formed by flattening the lower-triangular part of the attention matrix for each sentence at each layer, and N is the number of attention heads at that layer. The dependent variable (y) is a 7388-dimensional vector, where 7388 again reflects the length of the concatenated and flattened lower-triangular part of the BOLD response matrix for each sentence at each voxel for each subject. Since we constructed the BOLD matrix for each sentence based on the regressive eye saccade patterns, each BOLD matrix of each sentence is of size N_word_ × N_word_, its dimension matches the dimension of model’s attention matrix for each sentence.

The penalty regularization parameter for each voxel within each subject was determined using a grid search with nested cross-validation across 20 candidate regularization coefficients (log-spaced between 10 and 1,000), following previous approach (Huth et al., 2016). We adopted a train-test split method with 10-fold cross-validation, using 90% of the fMRI data (133 out of 148 sentences) to fit the ridge regression models and evaluating performance by computing the correlation between the predicted and observed time courses on the remaining 10% of the data (15 out of 148 sentences). For each voxel, a p-value for the correlation coefficient (r) was obtained by permuting the predicted time course 10,000 times and comparing the observed r to the distribution of permuted r values. of permuted r values. Significance of the r-map contrasts was assessed using cluster-based two sample t-tests with 10,000 permutations (Maris & Oostenveld, 2007). All our analyses and visualization were performed using custom python codes, making heavy use of the torch (v2.2.0), mne (v.1.6.1; Gramfort et al., 2014) and scipy (v1.12.0) packages.

While most prior model-brain alignment studies regressed embeddings at each model layer onto voxel-wise activity time series (e.g., Gao et al., 2024; Huth et al., 2016; Kumar et al., 2024; Schrimpf et al., 2021), this method is not feasible for the current study due to the non-linear nature of reading (Note that our task is not self-paced reading at word level where each word appears on the screen sequentially, instead, we presented the whole sentence on the screen and relied on eye-tracking to identify the timepoints for each word). We cannot directly regress the embeddings for each sentence with the fMRI data because is not strictly sequential. For example, our first participant read the sentence “Could humans live on Mars some day” in the following order based their eye fixations: “humans humans Could humans on Mars on some some day”. We could not simply input this disordered sequence into the LLM, as it would generate meaningless representations. Similarly, we cannot directly regress embeddings from the correctly ordered sentence onto the fMRI data, as the recorded neural responses correspond to the actual reading sequence, which does not follow the original sentence structure. To the best of our knowledge, no prior studies have employed this approach, likely because most previous research relied on naturalistic listening or self-paced reading paradigms at the word level, which inherently enforce sequential word processing. We hope our rationale is now clearer and that future studies incorporating concurrent eye-tracking and fMRI will consider applying our methods.

### Computational resources

Obtaining the model attention scores requires around 0.5 GPU hours for each model on a platform with 20 Intel Xeon Gold 6248 CPUs, 216 GB ROM, and 4 Nvidia Tesla v100 32 GB GPUs. Running each regression requires around 0.5 hours per model layer for each subject’s eye movement and fMRI data on a platform with 112 AMD EPYC 7522 CPUs and 512 GB ROM.

## Data availability

The attention matrices of all the LLMs for our experimental stimuli are available at https://github.com/RiverGao/scaling_finetuning. The eye-tracking and fMRI dataset is available at https://openneuro.org/datasets/ds003974/versions/3.0.0.

## Code availability

All codes are available at https://github.com/RiverGao/scaling_finetuning.

## Supplementary information

**Supplementary Table 1.** Mean and standard deviation of next-word prediction (NWP) loss from all LLMs averaged over all sentences in the test stimuli.

| Model | Mean | Standard deviation |
| --- | --- | --- |
| GPT-2 large | 5.268 | 1.671 |
| LLaMA 7B | 4.876 | 1.050 |
| Alpaca 7B | 5.378 | 1.196 |
| Vicuna 7B | 5.256 | 1.180 |
| Gemma-Instruct 7B | 5.947 | 1.701 |
| Mistral-Instruct 7B | 5.227 | 1.149 |
| LLaMA 13B | 4.873 | 1.078 |
| Alpaca 13B | 5.399 | 1.534 |
| Vicuna 13B | 5.781 | 1.320 |
| LLaMA 30B | 4.801 | 1.030 |
| LLaMA 65B | 4.805 | 1.067 |

**Supplementary Table 2.** Pairwise *t*-test results of the next-token prediction (NWP) loss from pairs of LLMs. Only pairs with a significant (p<0.05) or marginally significant (p<0.1) results are shown.

| model-pair | *t* | *p* |
| --- | --- | --- |
| GPT-2 large vs. LLaMA 7B | 8.688 | < 0.001 |
| GPT-2 large vs. Alpaca 7B | -3.819 | < 0.001 |
| GPT-2 large vs. Gemma-Instruct 7B | -7.398 | < 0.001 |
| GPT-2 large vs. LLaMA 13B | 8.370 | < 0.001 |
| GPT-2 large vs. Alpaca 13B | -2.327 | 0.026 |
| GPT-2 large vs. Vicuna 13B | -10.323 | < 0.001 |
| GPT-2 large vs. LLaMA 30B | 9.772 | < 0.001 |
| GPT-2 large vs. LLaMA 65B | 9.466 | < 0.001 |
| LLaMA 7B vs. Alpaca 7B | -17.695 | < 0.001 |
| LLaMA 7B vs. Vicuna 7B | -15.218 | < 0.001 |
| LLaMA 7B vs. Gemma-Instruct 7B | -13.322 | < 0.001 |
| LLaMA 7B vs. Mistral-Instruct 7B | -11.634 | < 0.001 |
| LLaMA 7B vs. Alpaca 13B | -6.439 | < 0.001 |
| LLaMA 7B vs. Vicuna 13B | -20.090 | < 0.001 |
| LLaMA 7B vs. LLaMA 30B | 3.589 | 0.001 |
| LLaMA 7B vs. LLaMA 65B | 2.659 | 0.011 |
| Alpaca 7B vs. Vicuna 7B | 4.731 | < 0.001 |
| Alpaca 7B vs. Gemma-Instruct 7B | -6.351 | < 0.001 |
| Alpaca 7B vs. Mistral-Instruct 7B | 3.208 | 0.002 |
| Alpaca 7B vs. LLaMA 13B | 15.111 | < 0.001 |
| Alpaca 7B vs. Vicuna 13B | -8.947 | < 0.001 |
| Alpaca 7B vs. LLaMA 30B | 17.186 | < 0.001 |
| Alpaca 7B vs. LLaMA 65B | 15.736 | < 0.001 |
| Vicuna 7B vs. Gemma-Instruct 7B | -7.897 | < 0.001 |
| Vicuna 7B vs. LLaMA 13B | 12.876 | < 0.001 |
| Vicuna 7B vs. Alpaca 13B | -1.806 | 0.085 |
| Vicuna 7B vs. Vicuna 13B | -12.652 | < 0.001 |
| Vicuna 7B vs. LLaMA 30B | 15.047 | < 0.001 |
| Vicuna 7B vs. LLaMA 65B | 14.424 | < 0.001 |
| Gemma-Instruct 7B vs. Mistral-Instruct 7B | 8.087 | < 0.001 |
| Gemma-Instruct 7B vs. LLaMA 13B | 13.417 | < 0.001 |
| Gemma-Instruct 7B vs. Alpaca 13B | 3.906 | < 0.001 |
| Gemma-Instruct 7B vs. LLaMA 30B | 14.926 | < 0.001 |
| Gemma-Instruct 7B vs. LLaMA 65B | 14.245 | < 0.001 |
| Mistral-Instruct 7B vs. LLaMA 13B | 12.935 | < 0.001 |
| Mistral-Instruct 7B vs. Alpaca 13B | -1.786 | 0.087 |
| Mistral-Instruct 7B vs. Vicuna 13B | -11.036 | < 0.001 |
| Mistral-Instruct 7B vs. LLaMA 30B | 15.733 | < 0.001 |
| Mistral-Instruct 7B vs. LLaMA 65B | 15.298 | < 0.001 |
| LLaMA 13B vs. Alpaca 13B | -6.290 | < 0.001 |
| LLaMA 13B vs. Vicuna 13B | -23.185 | < 0.001 |
| LLaMA 13B vs. LLaMA 30B | 4.521 | < 0.001 |
| LLaMA 13B vs. LLaMA 65B | 3.520 | 0.001 |
| Alpaca 13B vs. Vicuna 13B | -4.218 | < 0.001 |
| Alpaca 13B vs. LLaMA 30B | 7.171 | < 0.001 |
| Alpaca 13B vs. LLaMA 65B | 6.920 | < 0.001 |
| Vicuna 13B vs. LLaMA 30B | 22.438 | < 0.001 |
| Vicuna 13B vs. LLaMA 65B | 23.209 | < 0.001 |

**Supplementary Table 3.**
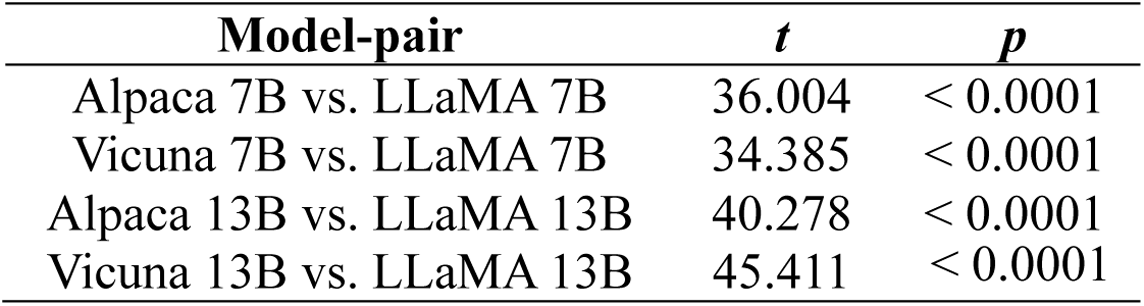
One sample *t*-test results for the divergence of the attention matrices for the experimental stimuli between the base and fine-tuned LLMs.]

| Model-pair | *t* | *p* |
| --- | --- | --- |
| Alpaca 7B vs. LLaMA 7B | 36.004 | < 0.0001 |
| Vicuna 7B vs. LLaMA 7B | 34.385 | < 0.0001 |
| Alpaca 13B vs. LLaMA 13B | 40.278 | < 0.0001 |
| Vicuna 13B vs. LLaMA 13B | 45.411 | < 0.0001 |

**Supplementary Table 4.** One sample *t*-test results for the divergence of the attention matrices for stimuli sentences with and without instruction or noise prefixes.

| Prefix | Model | *t* | *p* |
| --- | --- | --- | --- |
| instruction | LLaMA 7B | 40.837 | < 0.0001 |
|  | Alpaca 7B | 41.077 | < 0.0001 |
|  | Vicuna 7B | 43.583 | < 0.0001 |
|  | LLaMA 13B | 39.315 | < 0.0001 |
|  | Alpaca 13B | 44.100 | < 0.0001 |
|  | Vicuna 13B | 44.118 | < 0.0001 |
| noise | LLaMA 7B | 40.572 | < 0.0001 |
|  | Alpaca 7B | 40.732 | < 0.0001 |
|  | LLaMA 7B | 42.266 | < 0.0001 |
|  | LLaMA 13B | 37.969 | < 0.0001 |
|  | Alpaca 13B | 45.329 | < 0.0001 |
|  | Vicuna 13B | 47.389 | < 0.0001 |

**Supplementary Table 5.**
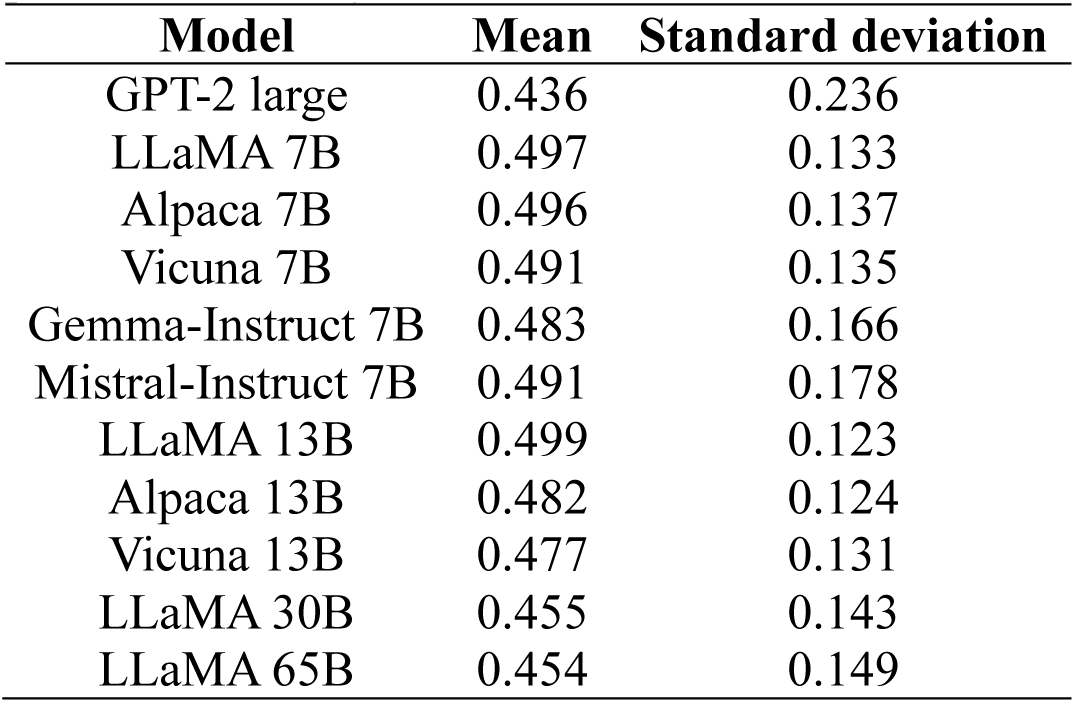
Mean regression scores of LLMs’ attention matrices against trivial patterns across layers.

**Supplementary Table 6.**
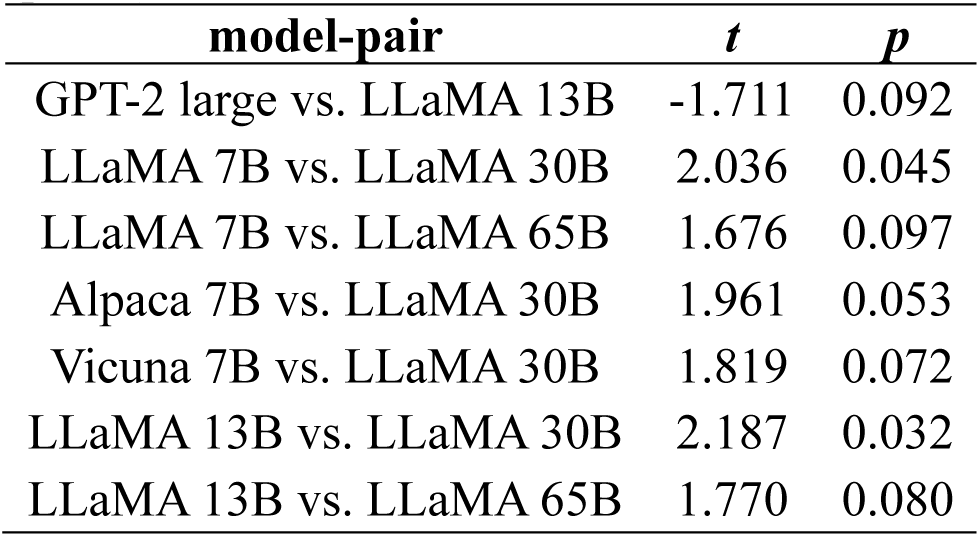
Pairwise t-test results for the mean regression scores of LLMs’ attention matrices against trivial patterns. Only pairs with a significant (p<0.05) or marginally significant (p<0.1) results are shown.

**Supplementary Table 7.** Mean regression scores from the best-performing layer of LLMs against eye regression numbers across participants.

| Model | Mean | Standard deviation |
| --- | --- | --- |
| GPT-2 base | 0.179 | 0.027 |
| GPT-2 medium | 0.186 | 0.028 |
| GPT-2 large | 0.174 | 0.026 |
| GPT-2 xlarge | 0.176 | 0.026 |
| LLaMA 7B | 0.346 | 0.032 |
| Alpaca 7B | 0.352 | 0.039 |
| Vicuna 7B | 0.344 | 0.034 |
| Gemma-Instruct 7B | 0.366 | 0.045 |
| Mistral-Instruct 7B | 0.377 | 0.047 |
| LLaMA 13B | 0.364 | 0.032 |
| Alpaca 13B | 0.374 | 0.035 |
| Vicuna 13B | 0.361 | 0.033 |
| LLaMA 30B | 0.387 | 0.038 |
| LLaMA 65B | 0.406 | 0.044 |

**Supplementary Table 8.** Pairwise t-test results for the mean regression scores from the best-performing layer of LLMs against eye regression numbers. Only pairs with a significant (p<0.05) or marginally significant (p<0.1) difference are shown.

| model-pair | *t* | *p* |
| --- | --- | --- |
| GPT-2 base vs. LLaMA 7B | -17.229 | < 0.001 |
| GPT-2 base vs. Alpaca 7B | -14.375 | < 0.001 |
| GPT-2 base vs. Vicuna 7B | -15.752 | < 0.001 |
| GPT-2 base vs. Gemma-Instruct 7B | -11.562 | < 0.001 |
| GPT-2 base vs. Mistral-Instruct 7B | -11.707 | < 0.001 |
| GPT-2 base vs. LLaMA 13B | -18.232 | < 0.001 |
| GPT-2 base vs. Alpaca 13B | -16.521 | < 0.001 |
| GPT-2 base vs. Vicuna 13B | -17.256 | < 0.001 |
| GPT-2 base vs. LLaMA 30B | -17.343 | < 0.001 |
| GPT-2 base vs. LLaMA 65B | -16.907 | < 0.001 |
| GPT-2 medium vs. LLaMA 7B | -21.790 | < 0.001 |
| GPT-2 medium vs. Alpaca 7B | -18.620 | < 0.001 |
| GPT-2 medium vs. Vicuna 7B | -20.070 | < 0.001 |
| GPT-2 medium vs. Gemma-Instruct 7B | -15.062 | < 0.001 |
| GPT-2 medium vs. Mistral-Instruct 7B | -15.394 | < 0.001 |
| GPT-2 medium vs. LLaMA 13B | -23.367 | < 0.001 |
| GPT-2 medium vs. Alpaca 13B | -21.375 | < 0.001 |
| GPT-2 medium vs. Vicuna 13B | -22.200 | < 0.001 |
| GPT-2 medium vs. LLaMA 30B | -23.003 | < 0.001 |
| GPT-2 medium vs. LLaMA 65B | -22.832 | < 0.001 |
| GPT-2 large vs. LLaMA 7B | -25.109 | < 0.001 |
| GPT-2 large vs. Alpaca 7B | -21.870 | < 0.001 |
| GPT-2 large vs. Vicuna 7B | -23.289 | < 0.001 |
| GPT-2 large vs. Gemma-Instruct 7B | -17.832 | < 0.001 |
| GPT-2 large vs. Mistral-Instruct 7B | -18.349 | < 0.001 |
| GPT-2 large vs. LLaMA 13B | -27.177 | < 0.001 |
| GPT-2 large vs. Alpaca 13B | -25.071 | < 0.001 |
| GPT-2 large vs. Vicuna 13B | -25.919 | < 0.001 |
| GPT-2 large vs. LLaMA 30B | -27.486 | < 0.001 |
| GPT-2 large vs. LLaMA 65B | -27.658 | < 0.001 |
| GPT-2 xlarge vs. LLaMA 7B | -28.541 | < 0.001 |
| GPT-2 xlarge vs. Alpaca 7B | -24.885 | < 0.001 |
| GPT-2 xlarge vs. Vicuna 7B | -26.470 | < 0.001 |
| GPT-2 xlarge vs. Gemma-Instruct 7B | -20.281 | < 0.001 |
| GPT-2 xlarge vs. Mistral-Instruct 7B | -20.884 | < 0.001 |
| GPT-2 xlarge vs. LLaMA 13B | -30.912 | < 0.001 |
| GPT-2 xlarge vs. Alpaca 13B | -28.524 | < 0.001 |
| GPT-2 xlarge vs. Vicuna 13B | -29.479 | < 0.001 |
| GPT-2 xlarge vs. LLaMA 30B | -31.325 | < 0.001 |
| GPT-2 xlarge vs. LLaMA 65B | -31.573 | < 0.001 |
| LLaMA 7B vs. LLaMA 30B | -2.052 | 0.070 |
| LLaMA 7B vs. LLaMA 65B | -3.961 | < 0.001 |
| Alpaca 7B vs. LLaMA 30B | -2.267 | 0.043 |
| Alpaca 7B vs. LLaMA 65B | -4.120 | < 0.001 |
| Vicuna 7B vs. LLaMA 30B | -2.876 | 0.009 |
| Vicuna 7B vs. LLaMA 65B | -4.719 | < 0.001 |
| Gemma 7B vs. LLaMA 13B | -2.592 | 0.021 |
| Gemma 7B vs. Alpaca 13B | -2.013 | 0.073 |
| Gemma 7B vs. Vicuna 13B | -2.068 | 0.070 |
| Gemma 7B vs. LLaMA 30B | -3.764 | 0.001 |
| Gemma 7B vs. LLaMA 65B | -5.415 | < 0.001 |
| Mistral-Instruct 7B vs. LLaMA 13B | -2.025 | 0.073 |
| Mistral-Instruct 7B vs. LLaMA 30B | -3.254 | 0.003 |
| Mistral-Instruct 7B vs. LLaMA 65B | -5.033 | < 0.001 |
| LLaMA 13B vs. LLaMA 65B | -3.597 | 0.001 |
| Alpaca 13B vs. LLaMA 30B | -2.011 | 0.073 |
| Alpaca 13B vs. LLaMA 65B | -4.110 | < 0.001 |
| Vicuna 13B vs. LLaMA 30B | -2.043 | 0.070 |
| Vicuna 13B vs. LLaMA 65B | -4.156 | < 0.001 |
| LLaMA 30B vs. LLaMA 65B | -2.475 | 0.025 |

**Supplementary Table 9.**
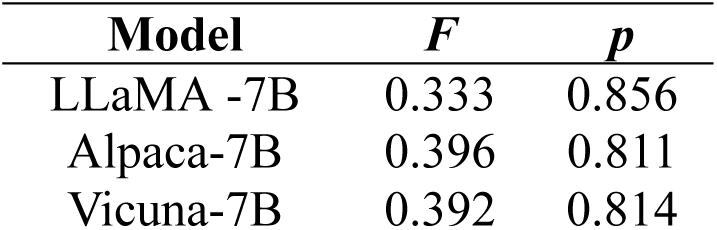
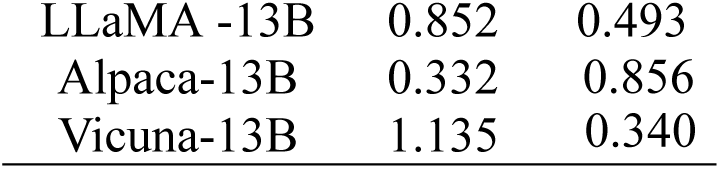
One-way ANOVA results for the regression scores of the attention matrices of each base and fine-tuned LLM against human regressive eye saccade number patterns for the stimuli sentences across the 5 experimental sections.

**Supplementary Table 10.** Mean normalized correlation coefficients of the ridge regression results from the best-performing layer of LLMs against fMRI data across participants.

| Model | Mean | Standard deviation |
| --- | --- | --- |
| GPT-2 base | 0.384 | 0.049 |
| GPT-2 medium | 0.434 | 0.059 |
| GPT-2 large | 0.458 | 0.063 |
| GPT-2 xlarge | 0.452 | 0.053 |
| LLaMA 7B | 0.500 | 0.070 |
| Alpaca 7B | 0.501 | 0.069 |
| Vicuna 7B | 0.492 | 0.070 |
| Gemma-Instruct 7B | 0.425 | 0.056 |
| Mistral-Instruct 7B | 0.471 | 0.058 |
| LLaMA 13B | 0.583 | 0.063 |
| Alpaca 13B | 0.579 | 0.063 |
| Vicuna 13B | 0.576 | 0.064 |
| LLaMA 30B | 0.586 | 0.072 |
| LLaMA 65B | 0.624 | 0.068 |

**Supplementary Table 11.** Pairwise t-test results for the mean correlation coefficients of the ridge regression results from the best-performing layer of LLMs against fMRI data. Only pairs with a significant (p<0.05) or marginally significant (p<0.1) difference are shown.

| model-pair | *t* | *p* |
| --- | --- | --- |
| GPT-2 medium vs. GPT-2 base | 4.693 | <0.001 |
| GPT-2 large vs. GPT-2 base | 3.309 | 0.003 |
| GPT-2 xlarge vs. GPT-2 base | 7.764 | <0.001 |
| LLaMA 7B vs. GPT-2 base | 5.829 | <0.001 |
| Alpaca 7B vs. GPT-2 base | 6.886 | <0.001 |
| Vicuna 7B vs. GPT-2 base | 5.818 | <0.001 |
| Gemma-Instruct 7B vs. GPT-2 base | 2.918 | 0.009 |
| Mistral-Instruct 7B vs. GPT-2 base | 5.065 | <0.001 |
| LLaMA 13B vs. GPT-2 base | 14.439 | <0.001 |
| Alpaca 13B vs. GPT-2 base | 12.045 | <0.001 |
| Vicuna 13B vs. GPT-2 base | 12.176 | <0.001 |
| LLaMA 30B vs. GPT-2 base | 16.533 | <0.001 |
| LLaMA 65B vs. GPT-2 base | 15.235 | <0.001 |
| GPT-2 xlarge vs. GPT-2 medium | 1.978 | 0.073 |
| LLaMA 7B vs. GPT-2 medium | 1.842 | 0.095 |
| Alpaca 7B vs. GPT-2 medium | 2.392 | 0.032 |
| LLaMA 13B vs. GPT-2 medium | 7.48 | <0.001 |
| Alpaca 13B vs. GPT-2 medium | 6.104 | <0.001 |
| Vicuna 13B vs. GPT-2 medium | 6.36 | <0.001 |
| LLaMA 30B vs. GPT-2 medium | 8.303 | <0.001 |
| LLaMA 65B vs. GPT-2 medium | 9.136 | <0.001 |
| GPT-2 xlarge vs. GPT-2 large | 2.522 | 0.023 |
| LLaMA 7B vs. GPT-2 large | 2.322 | 0.036 |
| Alpaca 7B vs. GPT-2 large | 2.877 | 0.01 |
| Vicuna 7B vs. GPT-2 large | 2.141 | 0.054 |
| LLaMA 13B vs. GPT-2 large | 7.409 | <0.001 |
| Alpaca 13B vs. GPT-2 large | 6.258 | <0.001 |
| Vicuna 13B vs. GPT-2 large | 6.54 | <0.001 |
| LLaMA 30B vs. GPT-2 large | 7.987 | <0.001 |
| LLaMA 65B vs. GPT-2 large | 9.159 | <0.001 |
| GPT-2 xlarge vs. Gemma-Instruct 7B | 3.68 | 0.001 |
| GPT-2 xlarge vs. Mistral-Instruct 7B | 2.049 | 0.062 |
| LLaMA 13B vs. GPT-2 xlarge | 6.302 | <0.001 |
| Alpaca 13B vs. GPT-2 xlarge | 4.844 | <0.001 |
| Vicuna 13B vs. GPT-2 xlarge | 5.133 | <0.001 |
| LLaMA 30B vs. GPT-2 xlarge | 7.144 | <0.001 |
| LLaMA 65B vs. GPT-2 xlarge | 8.256 | <0.001 |
| LLaMA 7B vs. Gemma-Instruct 7B | 3.175 | 0.005 |
| LLaMA 7B vs. Mistral-Instruct 7B | 1.94 | 0.077 |
| LLaMA 13B vs. LLaMA 7B | 4.233 | <0.001 |
| Alpaca 13B vs. LLaMA 7B | 3.287 | 0.004 |
| Vicuna 13B vs. LLaMA 7B | 3.551 | 0.002 |
| LLaMA 30B vs. LLaMA 7B | 4.641 | <0.001 |
| LLaMA 65B vs. LLaMA 7B | 6.012 | <0.001 |
| Alpaca 7B vs. Gemma-Instruct 7B | 3.857 | 0.001 |
| Alpaca 7B vs. Mistral-Instruct 7B | 2.527 | 0.023 |
| LLaMA 13B vs. Alpaca 7B | 4.265 | <0.001 |
| Alpaca 13B vs. Alpaca 7B | 3.223 | 0.004 |
| Vicuna 13B vs. Alpaca 7B | 3.504 | 0.002 |
| LLaMA 30B vs. Alpaca 7B | 4.728 | <0.001 |
| LLaMA 65B vs. Alpaca 7B | 6.186 | <0.001 |
| Vicuna 7B vs. Gemma-Instruct 7B | 3.012 | 0.007 |
| LLaMA 13B vs. Vicuna 7B | 4.63 | <0.001 |
| Alpaca 13B vs. Vicuna 7B | 3.62 | 0.002 |
| Vicuna 13B vs. Vicuna 7B | 3.881 | 0.001 |
| LLaMA 30B vs. Vicuna 7B | 5.102 | <0.001 |
| LLaMA 65B vs. Vicuna 7B | 6.384 | <0.001 |
| LLaMA 13B vs. Gemma-Instruct 7B | 9.174 | <0.002 |
| Alpaca 13B vs. Gemma-Instruct 7B | 7.747 | <0.003 |
| Vicuna 13B vs. Gemma-Instruct 7B | 8.021 | <0.004 |
| LLaMA 30B vs. Gemma-Instruct 7B | 10.012 | <0.005 |
| LLaMA 65B vs. Gemma-Instruct 7B | 10.87 | <0.006 |
| LLaMA 13B vs. Mistral-Instruct 7B | 7.943 | <0.007 |
| Alpaca 13B vs. Mistral-Instruct 7B | 6.5 | <0.008 |
| Vicuna 13B vs. Mistral-Instruct 7B | 6.786 | <0.009 |
| LLaMA 30B vs. Mistral-Instruct 7B | 8.772 | <0.010 |
| LLaMA 65B vs. Mistral-Instruct 7B | 9.793 | <0.011 |
| LLaMA 65B vs. LLaMA 13B | 2.363 | 0.034 |
| LLaMA 65B vs. Alpaca 13B | 3.217 | 0.004 |
| LLaMA 65B vs. Vicuna 13B | 2.921 | 0.009 |
| LLaMA 65B vs. LLaMA 30B | 2.127 | 0.055 |

### Additional results from fMRI data of naturalistic listening

To verify whether our findings can generalize to a different dataset, we performed the same analysis on a fMRI dataset collected while participants listened to a 20-minute Chinese audiobook in the scanner. The dataset was collected for another project(Fischl, 2012)(Wang et al., 2025) and involves a total of 26 participants (15 females, mean age=23.96±2.23 years) listening to two sections of the Chinese version of “The Little Prince”. All participants were right-handed native Mandarin speakers enrolled in an undergraduate or graduate program in Shanghai and had no self-reported history of neurological disorders. The fMRI data was collected in a 7.0 T Terra Siemens MRI scanner at the Zhangjiang International Brain Imaging Centre at Fudan University, Shanghai. Anatomical scans were obtained using a Magnetization Prepared RApid Gradient-Echo (MP-RAGE) SAG iPAT2 pulse sequence with T1-weighted contrast (256 single-shot interleaved sagittal slices with A/P phase encoding direction; voxel size: 0.7×0.7×0.7 mm; FOV: 208 mm; TR: 3800 ms; TE: 2.32 ms; flip angle: 7°; acquisition time: 3 s; GRAPPA in-plane acceleration factor: 3). Functional imaging was conducted with T2-weighted echo-planar imaging (85 interleaved axial slices, anterior-posterior phase encoding; voxel size: 1.6×1.6×1.6 mm; FOV: 208 mm; TR: 1000 ms; TE: 22.2 ms; multiband acceleration factor: 5; flip angle: 45°). fMRI preprocessing of the was conducted using fMRIPrep (v25.0.0) with all default parameters. Anatomical images were corrected for intensity non-uniformity (N4BiasFieldCorrection), skull-stripped using ANTs-based extraction (OASIS30ANTs template), and segmented into tissue classes using FSL’s ‘fast’. Brain surfaces were reconstructed using ‘recon-all’ (FreeSurfer 7.3.2).

We regressed the attention weights of the base and fine-tuned LLaMA3 8B and LLaMA3 70B models against fMRI data matrices at the paragraph level. This analysis extends beyond sentence-level comprehension to discourse-level processing and incorporates both a different modality (listening vs. reading) and a different language (Chinese vs. English). Specifically, we extracted the BOLD signal time-locked to each word’s offset, adding five scans to account for peak hemodynamic responses within each paragraph of the stimuli. We then constructed a representational dissimilarity matrix (RDM; Kriegeskorte et al., 2008) for each paragraph using the 20 neighboring voxels around each voxel. The lower triangles of these RDMs were extracted, concatenated across all paragraphs, and used as dependent variables. For our predictors, we extracted and flattened the lower triangles from the attention matrices for all paragraphs. We then applied the same ridge regression analyses using attention matrices from all attention heads to predict the fMRI data RDM at each voxel within a bilateral language mask. The language mask covered regions including the whole temporal lobe, the inferior frontal gyrus (IFG) and the angular gyrus (AG).

Our findings remained consistent: Model scaling had a significant effect on model-brain alignment, while fine-tuned and base models of the same size showed no difference in brain encoding performance (see Supplementary Fig. 1a and Supplementary Table 12-14).

### Additional results from fMRI data of comprehension tasks

To further investigate the impact of fine-tuning on model-brain alignment in a task-intensive setting, we regressed predictions from LLaMA3-Instruct (8B and 70B) against fMRI data while participants answered multiple-choice comprehension questions about the preceding listening session through button press in the scanner. The full set of questions is available in our OpenNeuro dataset directory (https://openneuro.org/datasets/ds005345). An example question (translated into English) is as follows:

Why is it difficult to talk to the Little Prince?

a. He doesn’t speak.
b. He doesn’t ask questions.
c. He speaks an alien language.
d. He doesn’t answer questions directly.

We prompted the fine-tuned LLMs with the text from the listening session along with the multiple-choice questions. We then extracted the models’ probability for the correct answer and computed its surprisal (negative log probability) to regress against the fMRI scans time-locked to the button press for each question (adding 5 scans to capture the peak hemodynamic response). The regression was conducted for each voxel within a bilateral language mask in every subject. At the group level, the beta maps for each model were z-scored and assessed for statistical significance using cluster-based permutation t-tests (Maris & Oostenveld, 2007) with 10,000 permutations.

Our results indicated that for smaller model sizes, LLaMA3 exhibited higher mean regression scores with task-based fMRI data compared to LLaMA3-Instruct. For the larger model size, LLaMA3-Instruct had a higher mean regression score across participants relative to the base model (see Supplementary Fig. 1b and Supplementary Table 15-16).

**Supplementary Fig. 1.**
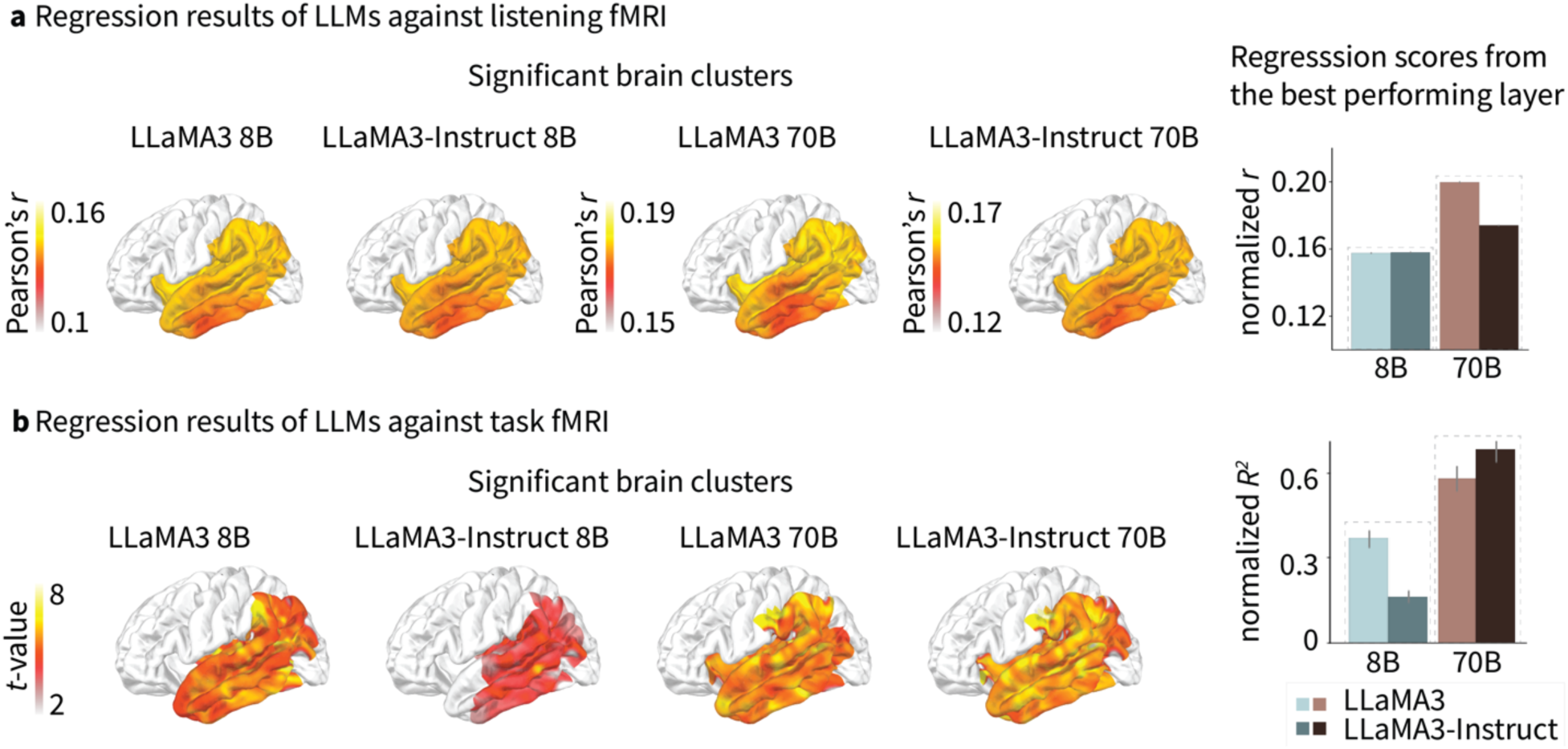
Regression results from additional fMRI datasets. **a,** Significant brain clusters showing alignment between LLMs and fMRI data during naturalistic listening. The right panel displays the normalized Pearson’s *r* averaged over the significant brain clusters from the best-performing layer across participants. **b,** Significant brain clusters showing alignment between LLMs and fMRI data during the question-answering task. The right panel presents the averaged regression scores from the best-performing layer across participants.

**Supplementary Table 12.**
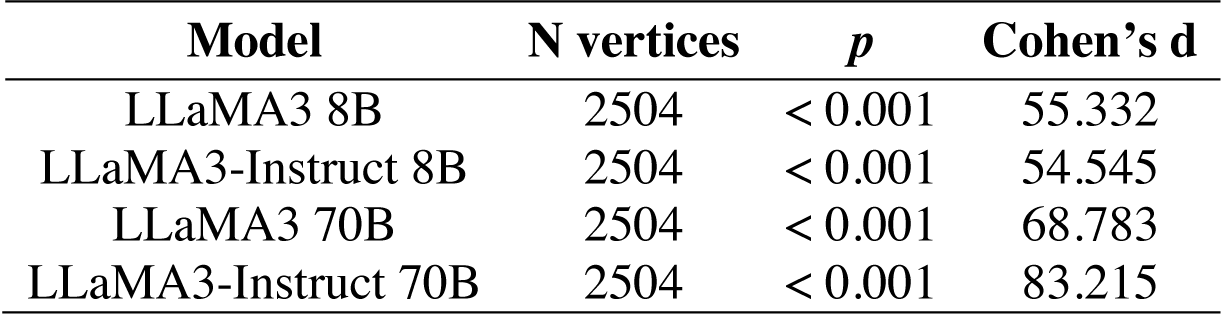
Summary statistics for the significant brain clusters from the correlation coefficients of LLMs against listening fMRI.

| Model | N vertices | $p$ | Cohen's $d$ |
| --- | --- | --- | --- |
| LLaMA3 8B | 2504 | < 0.001 | 55.332 |
| LLaMA3-Instruct 8B | 2504 | < 0.001 | 54.545 |
| LLaMA3 70B | 2504 | < 0.001 | 68.783 |
| LLaMA3-Instruct 70B | 2504 | < 0.001 | 83.215 |

**Supplementary Table 13.** Mean normalized correlation coefficients of the best-performing layer of LLMs against listening fMRI across participants.

| Model | Mean | Standard deviation |
| --- | --- | --- |
| LLaMA3 8B | 0.158 | 0.003 |
| LLaMA3-Instruct 8B | 0.158 | 0.003 |
| LLaMA3 70B | 0.199 | 0.003 |
| LLaMA3-Instruct 70B | 0.174 | 0.002 |

**Supplementary Table 14.** Pairwise t-test results for the correlation coefficients from the best-performing layer of LLMs against listening fMRI data. Only pairs with a significant (p<0.05) or marginally significant (p<0.1) difference are shown.

| Model-pair | $t$ | $p$ |
| --- | --- | --- |
| LLaMA3-Instruct 8B vs. LLaMA3 8B | 21.173 | < 0.001 |
| LLaMA3 70B vs. LLaMA3 8B | 2561.722 | < 0.001 |
| LLaMA3-Instruct 70B vs. LLaMA3 8B | 84.477 | < 0.001 |
| LLaMA3 70B vs. LLaMA3-Instruct 8B | 1824.727 | < 0.001 |
| LLaMA3-Instruct 70B vs. LLaMA3-Instruct 8B | 84.885 | < 0.001 |
| LLaMA3 70B vs. LLaMA3-Instruct 70B | 122.611 | < 0.001 |

**Supplementary Table 15.** Summary statistics for the significant brain clusters from the beta values of LLMs against task fMRI.

| Model | N vertices | *p* | *t* | Cohen's *d* |
| --- | --- | --- | --- | --- |
| LLaMA3 8B | 1094 | < 0.001 | 5.410 | 7.805 |
| LLaMA3-Instruct 8B | 705 | < 0.001 | 4.323 | 7.366 |
| LLaMA3 70B | 1439 | < 0.001 | 5.557 | 9.814 |
| LLaMA3-Instruct 70B | 1652 | < 0.001 | 5.664 | 8.473 |

**Supplementary Table 16.** Mean normalized regression scores of the best-performing layer of LLMs against task fMRI across participants.

| Model | Mean | Standard deviation |
| --- | --- | --- |
| LLaMA3 8B | 0.365 | 0.153 |
| LLaMA3-Instruct 8B | 0.162 | 0.108 |
| LLaMA3 70B | 0.578 | 0.231 |
| LLaMA3-Instruct 70B | 0.682 | 0.221 |

**Supplementary Table 17.** Pairwise t-test results for the beta maps of the regression results from the best-performing layer of LLMs against listening fMRI data. Only pairs with a significant (p<0.05) or marginally significant (p<0.1) difference are shown.

| Model-pair | *t* | *p* |
| --- | --- | --- |
| LLaMA3 8B vs. LLaMA3-Instruct 8B | 6.874 | < 0.001 |
| LLaMA3 70B vs. LLaMA3 8B | 3.729 | < 0.001 |
| LLaMA3-Instruct 70B vs. LLaMA3 8B | 7.456 | < 0.001 |
| LLaMA3 70B vs. LLaMA3-Instruct 8B | 19.895 | < 0.001 |
| LLaMA3-Instruct 70B vs. LLaMA3-Instruct 8B | 39.124 | < 0.001 |
| LLaMA3-Instruct 70B vs. LLaMA3 70B | 4.083 | < 0.001 |

**Supplementary Fig. 2.**
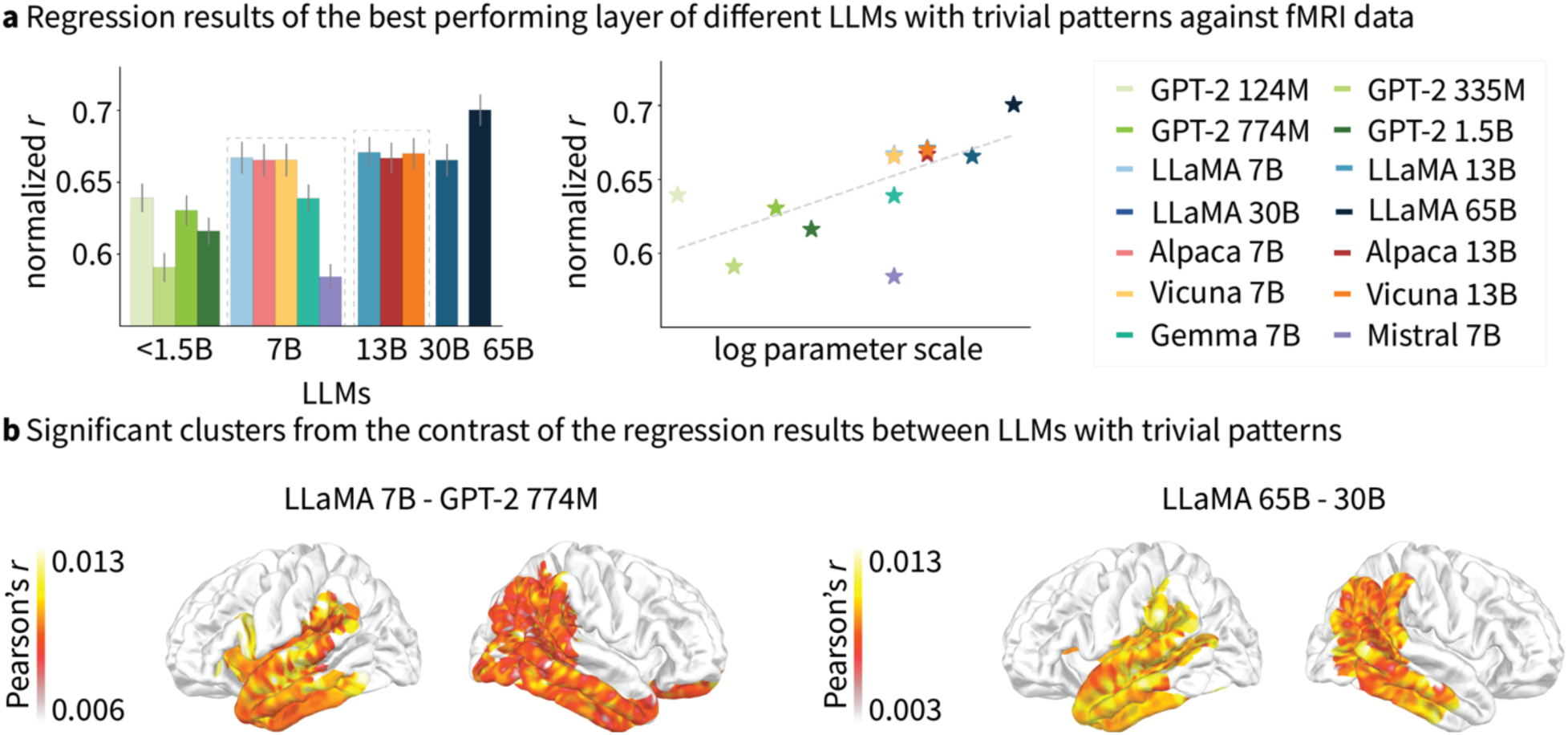
Regression results using original attention matrices of the LLMs (without subtracting the trivial patterns). **a,** Regression results of the LLMs’ best-performing layer on the fMRI activity patterns and their results on a logarithmic size scale. Error bars denote standard deviation. **b,** Significant brain clusters from the contrast of the correlation coefficients of different LLMs with smaller and larger sizes.

**Supplementary Table 18.** Mean correlation coefficients from the best-performing layer of LLMs using their original attention matrices against fMRI data across participants.

| Model | Mean | Standard deviation |
| --- | --- | --- |
| GPT-2 base | 0.639 | 0.069 |
| GPT-2 medium | 0.591 | 0.071 |
| GPT-2 large | 0.630 | 0.075 |
| GPT-2 xlarge | 0.616 | 0.067 |
| LLaMA 7B | 0.667 | 0.079 |
| Alpaca 7B | 0.665 | 0.080 |
| Vicuna 7B | 0.666 | 0.079 |
| Gemma-Instruct 7B | 0.639 | 0.069 |
| Mistral-Instruct 7B | 0.584 | 0.063 |
| LLaMA 13B | 0.671 | 0.076 |
| Alpaca 13B | 0.667 | 0.076 |
| Vicuna 13B | 0.670 | 0.076 |
| LLaMA 30B | 0.665 | 0.079 |
| LLaMA 65B | 0.700 | 0.077 |

**Supplementary Table 19.** Pairwise t-test results for the mean correlation coefficients from the best-performing layer of LLMs using their original attention matrices against fMRI data. Only pairs with a significant (p<0.05) or marginally significant (p<0.1) difference are shown.

| Model-pair | *t* | *p* |
| --- | --- | --- |
| GPT-2 base vs. GPT-2 medium | 2.915 | 0.027 |
| GPT-2 base vs. GPT-2 xlarge | 3.134 | 0.035 |
| LLaMA 7B vs. GPT-2 base | 2.283 | 0.071 |
| Alpaca 7B vs. GPT-2 base | 4.261 | 0.008 |
| Vicuna 7B vs. GPT-2 base | 4.720 | 0.009 |
| GPT-2 base vs. Mistral-Instruct 7B | 3.344 | 0.020 |
| LLaMA 13B vs. GPT-2 base | 2.235 | 0.089 |
| Vicuna 13B vs. GPT-2 base | 2.143 | 0.076 |
| LLaMA 30B vs. GPT-2 base | 4.980 | 0.001 |
| LLaMA 65B vs. GPT-2 base | 3.918 | 0.008 |
| GPT-2 large vs. GPT-2 medium | 3.168 | 0.008 |
| LLaMA 7B vs. GPT-2 medium | 4.653 | 0.002 |
| Alpaca 7B vs. GPT-2 medium | 6.192 | <0.001 |
| Vicuna 7B vs. GPT-2 medium | 5.182 | 0.002 |
| Gemma-Instruct 7B vs. GPT-2 medium | 3.404 | 0.011 |
| LLaMA 13B vs. GPT-2 medium | 4.256 | 0.005 |
| Alpaca 13B vs. GPT-2 medium | 4.062 | 0.005 |
| Vicuna 13B vs. GPT-2 medium | 4.929 | 0.001 |
| LLaMA 30B vs. GPT-2 medium | 8.538 | <0.001 |
| LLaMA 65B vs. GPT-2 medium | 6.591 | <0.001 |
| GPT-2 large vs. GPT-2 xlarge | 2.094 | 0.063 |
| LLaMA 7B vs. GPT-2 large | 2.361 | 0.038 |
| Alpaca 7B vs. GPT-2 large | 3.083 | 0.010 |
| Vicuna 7B vs. GPT-2 large | 2.072 | 0.065 |
| GPT-2 large vs. Mistral-Instruct 7B | 3.249 | 0.008 |
| LLaMA 30B vs. GPT-2 large | 4.334 | 0.001 |
| LLaMA 65B vs. GPT-2 large | 4.287 | 0.001 |
| LLaMA 7B vs. GPT-2 xlarge | 3.731 | 0.014 |
| Alpaca 7B vs. GPT-2 xlarge | 5.942 | 0.002 |
| Vicuna 7B vs. GPT-2 xlarge | 5.846 | 0.004 |
| Gemma 7B vs. GPT-2 xlarge | 2.597 | 0.048 |
| LLaMA 13B vs. GPT-2 xlarge | 3.775 | 0.020 |
| Alpaca 13B vs. GPT-2 xlarge | 3.597 | 0.016 |
| Vicuna 13B vs. GPT-2 xlarge | 4.349 | 0.005 |
| LLaMA 30B vs. GPT-2 xlarge | 7.535 | <0.001 |
| LLaMA 65B vs. GPT-2 xlarge | 5.404 | 0.002 |
| LLaMA 7B vs. Mistral 7B | 4.636 | 0.004 |
| Alpaca 7B vs. Mistral 7B | 6.422 | 0.001 |
| Alpaca 7B vs. Alpaca 13B | 2.342 | 0.058 |
| Alpaca 7B vs. Vicuna 13B | 2.006 | 0.085 |
| Vicuna 7B vs. Mistral 7B | 5.594 | 0.003 |
| Gemma-Instruct 7B vs. Mistral-Instruct 7B | 3.469 | 0.013 |
| LLaMA 30B vs. Gemma-Instruct 7B | 2.352 | 0.038 |
| LLaMA 65B vs. Gemma-Instruct 7B | 2.538 | 0.039 |
| LLaMA 13B vs. Mistral-Instruct 7B | 4.393 | 0.007 |
| Alpaca 13B vs. Mistral-Instruct 7B | 4.311 | 0.005 |
| Vicuna 13B vs. Mistral-Instruct 7B | 5.173 | 0.001 |
| LLaMA 30B vs. Mistral-Instruct 7B | 8.724 | <0.001 |
| LLaMA 65B vs. Mistral-Instruct 7B | 6.471 | <0.001 |
| LLaMA 30B vs. Alpaca 13B | 2.998 | 0.012 |
| LLaMA 65B vs. Alpaca 13B | 3.017 | 0.019 |
| LLaMA 30B vs. Vicuna 13B | 2.552 | 0.025 |
| LLaMA 65B vs. Vicuna 13B | 2.912 | 0.020 |
| LLaMA 65B vs. LLaMA 30B | 1.785 | 0.100 |

